# Integrity of the short arm of nuclear pore Y-complex is required for mouse embryonic stem cell growth and differentiation

**DOI:** 10.1101/2020.10.27.357376

**Authors:** Alba Gonzalez-Estevez, Annalisa Verrico, Clarisse Orniacki, Bernardo Reina-San-Martin, Valérie Doye

## Abstract

Many cellular processes, ranging from cell division to differentiation, are controlled by nuclear pore complexes (NPCs). However studying contributions of individual NPC subunits to these processes in vertebrates has long been impeded by their complexity and the lack of efficient genetic tools. Here we use genome editing in mouse embryonic stem cells (mESCs) to characterize the role of NPC structural components, focusing on the short arm of the Y-complex that comprises Nup85, Seh1 and Nup43. We show that Seh1 and Nup43, although dispensable in pluripotent mESCs, are required for their normal cell growth rates, their viability upon differentiation, and for the maintenance of proper NPC density. mESCs with an N-terminally truncated Nup85 mutation (in which interaction with Seh1 is greatly impaired) feature a similar reduction of NPC density. However, their proliferation and differentiation are unaltered, indicating that it is the integrity of the Y-complex, rather than the number of NPCs, that is critical to ensure these processes.

**Summary statement:** Seh1 and Nup43, although dispensable in pluripotent mouse embryonic stem cells, are required for normal cell growth, viability upon differentiation, and maintenance of proper NPC density.

## INTRODUCTION

Nuclear pore complexes (NPCs) are huge structures embedded in the nuclear envelope (NE). They provide the sole gateways for bidirectional nucleocytoplasmic transport, but also participate in a wide variety of other cellular processes including cell division and gene regulation (reviewed in Buchwalter et al., 2019; Hezwani and Fahrenkrog, 2017). NPCs are composed of ∼ 30 distinct proteins (called nucleoporins or Nups), each present in multiple copies and forming a ring with an eightfold rotational symmetry. Among them, structural Nups assemble to form a scaffold that anchors Nups with unfolded domains, cytoplasmic filaments and the nuclear basket (reviewed in Hampoelz et al., 2019; Lin and Hoelz, 2019).

The three-dimensional organization of the NPC scaffold has been determined at atomic resolution (reviewed in (Hampoelz et al., 2019; Lin and Hoelz, 2019). It is formed by an inner rim sandwiched by two outer (cytoplasmic and nuclear) rims, whose main component is the evolutionarily-conserved Y-complex. In metazoans, this complex (also named Nup107-160 complex) comprises Nup133, Nup107, Nup96 and Sec13 (forming the stem of the Y); Nup160, Nup37 and Elys (building the long arm); and Nup85, Seh1 (also named Seh1l) and Nup43 (forming the short arm)(**Fig. 1A**) (Loiodice et al., 2004; Rasala et al., 2006; von Appen et al., 2015).

**Figure 1:**
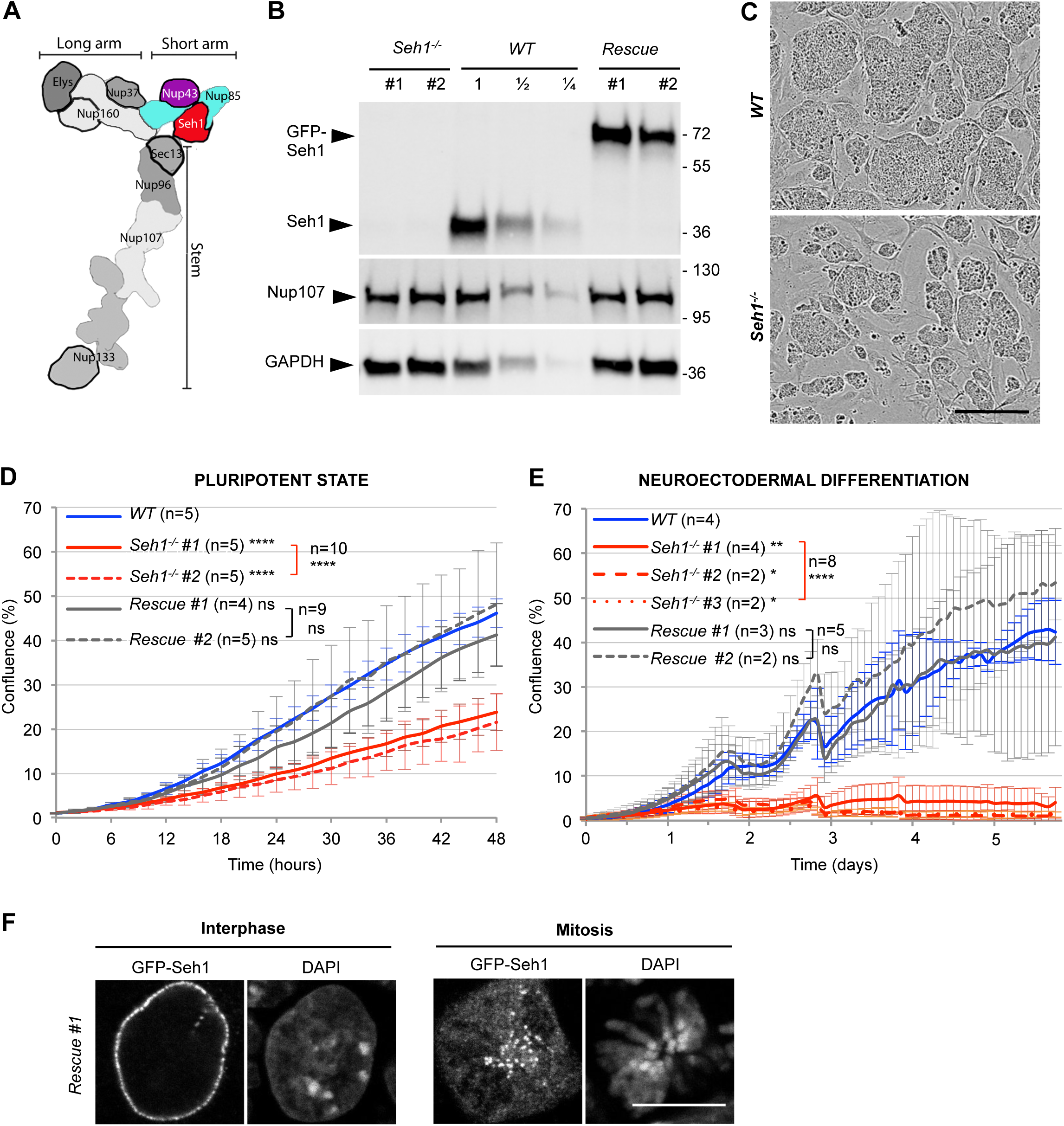
Seh1 depletion leads to cell growth delay and impaired cell survival upon differentiation. **A.** Schematic representation of the Y-complex (adapted from von Appen et al., 2015), highlighting the components of the short arm, namely Nup43 (shown in purple), Nup85 (blue) and Seh1 (red). ß-propellers are outlined with black strokes. **B.** Western-blot showing the expression of endogenous or GFP-tagged Seh1, Nup107 and GAPDH (used as loading control) in whole cell extracts from the indicated cell lines. 1/2 and 1/4 dilutions of *WT* extracts were also loaded. Molecular weights are indicated (kilodaltons). **C.** Representative phase contrast images of *WT* and *Seh1^-/-^* (#1) mESCs colonies acquired after 2 days of growth on the IncuCyte→ imager. Scale bar, 300µm. **D-E.** Confluence of *WT* (blue), *Seh1^-/-^* (#1 and #2, red) and *Rescue* (#1 and #2, grey) mESCs was quantified with the IncuCyte→ system either at pluripotent stage (**D**) or upon differentiation towards neuroectodermal lineage (**E**). Error bars correspond to the standard deviation arising from the indicated number of independent experiments [n]. Statistical analyses of these confluence curves were performed at the last time-points. Brackets indicate statistics performed using all values from cell lines bearing a given mutation, compared to *WT* (see Materials and Methods). **F.** Representative spinning disk images of *Rescue* (#1) mESCs showing proper localization of GFP-Seh1 at the NE in interphase (one plane) and at kinetochores in mitosis (a projection of 3 optical sections is presented). Scale bars, 10 µm.

Functional studies in vertebrates have shown that the Y-complex is critical for NPC assembly, both after mitosis and during interphase (Doucet et al., 2010; Harel et al., 2003; Walther et al., 2003). Studies in mammalian cells also showed that in mitosis a fraction of the Y-complex localizes at kinetochores (Loiodice et al., 2004; Rasala et al., 2006) where it is required for proper chromosome congression and segregation (Platani et al., 2009; Zuccolo et al., 2007). Because the members of the Y-complex (Y-Nups) are tightly associated throughout the cell cycle (Loiodice et al., 2004; Rabut et al., 2004), they were long anticipated to work as an entity. However, one of its components, Sec13, is also part of the COPII coat complex involved in vesicle budding (Salama et al., 1993). In addition, Sec13 and Seh1 also belong to the unrelated GATOR2 complex, an indirect regulator of the mTORC1 pathway that controls cell growth and proliferation (Bar-Peled et al., 2013), further complicating the study of their function in the context of the Y-complex.

In mice, inactivation of most Y-Nups genes (namely *Elys*, *Nup96*, *Nup133*, *Nup85*, *Sec13* and *Seh1*, but not *Nup37)* lead to embryonic lethality (Faria et al., 2006; Liu et al., 2019; Lupu et al., 2008; Moreira et al., 2015; Okita et al., 2004; Terashima et al., 2020; https://www.mousephenotype.org/data/genes/MGI:1919964). In particular, Nup133 was found to be essential for mouse development beyond gastrulation (Lupu et al., 2008). Studies performed in mouse embryonic stem cell (mESCs) showed that Nup133 is dispensable for cell growth at the pluripotent stage, but is required for mESC differentiation (Lupu et al., 2008). In mESCs, Nup133 is dispensable for NPC scaffold assembly but required for the proper assembly of the nuclear pore basket (Souquet et al., 2018). However, it is not clear if the role of Nup133 in NPC basket assembly underlies its functions in cell differentiation. More recently, Seh1, which is critical for proper mitotic progression in cancer cell lines (Platani et al., 2018; Platani et al., 2009; Zuccolo et al., 2007), was found to be required for differentiation of oligodendrocyte progenitors (Liu et al., 2019). However, the potential contribution of Seh1 to cell cycle progression in non-transformed cells and at other stages of cell differentiation needed to be addressed.

Here we assessed the requirements for Seh1 in pluripotent mESCs and upon their differentiation towards neuroectodermal lineage, determined whether these requirements reflect its role in the GATOR2-complex or in the short arm of the Y-complex, and further addressed the specific function of these proteins in NPC integrity. This systematic analysis enabled us to disentangle the processes underlying the contribution of these of these Y-Nups in NPC assembly, nuclear size, cell growth and differentiation.

## Results

### Seh1 is required for mESC growth and survival upon differentiation

Using CRISPR/Cas9 gene-editing technology we obtained several independent *Seh1^-/-^* mESC clones (of which three were further examined in this study; see Materials and Methods and Table S2) (**Fig. 1B**). This indicates that Seh1 is dispensable for mESC viability at the pluripotent stage. We noticed however that *Seh1*^-*/-*^ mESCs formed smaller colonies than did *WT* mESCs (**Fig. 1C**). Consistently, automated cell growth analyses of *Seh1^-/-^* mESCs showed a clear reduction of cell confluence compared to *WT* (**Fig. 1D**). More strikingly, *Seh1^-/-^* cells showed a strong impairment in viability from the very early stages of monolayer differentiation towards neuroectodermal lineage and almost no cells were recovered after 5 days (**Fig. 1E and Movies S1-S2**).

To verify the specificity of these phenotypes, we next integrated at the permissive *Tigre* locus (<u>Tig</u>htly <u>re</u>gulated; Zeng et al., 2008) of *Seh1^-/-^* mESCs a GFP-tagged *Seh1* cDNA expressed under the control of the pCAG promoter. The resulting cell lines (subsequently named “*Rescue”* #1 and #2) expressed GFP-Seh1 at a level comparable to that of the endogenous untagged protein (**Fig. 1B**). We observed a specific enrichment of GFP-Seh1 at nuclear pores in interphase and at kinetochores throughout mitosis (**Fig. 1F**). Most importantly, the growth rate of the *Rescue* cell lines was comparable to that of *WT* cells both at the pluripotent stage (**Fig. 1D**) and upon neuroectodermal differentiation (**Fig. 1E and Movie S3**).

To exclude the possibility that phenotypes observed in *Seh1^-/-^* mESCs at the pluripotent stage could be due to cell adaptation, we also generated cell lines in which endogenous *Seh1* was N-terminally tagged with the 7 kDa mini Auxin Inducible Degron (mAID) sequence, to induce its acute degradation upon auxin addition (Natsume et al., 2016). A GFP tag was also introduced to allow visualization of both the localization and degradation of the resulting GFP-mAID-Seh1 fusion. Upon addition of auxin to *GFP-mAID-Seh1* mESCs, the GFP signal rapidly declined in mitotic cells whereas, as also previously observed in HCT116 cells (Platani et al., 2018), the decay was more progressive in interphasic cells (**Fig. S1 A-D**). While the *GFP-mAID-Seh1* clones showed normal cell growth and differentiation properties in control conditions, addition of auxin recapitulated both the cell growth and differentiation defects observed in *Seh1^-/-^* mESCs (**Fig. S1 E, F)**.

Together these data reveal that the lack of Seh1 specifically causes an impaired cell growth of mESCs at the pluripotent stage and drastically reduced viability upon induction of neuroectodermal differentiation.

### The altered growth rate of pluripotent *Seh1^-/-^* mESCs is mainly caused by extended interphases

Seh1 is known to play a role in mitosis in cancer cell lines, in which its depletion causes a delay in mitotic progression associated with chromosome congression and segregation defects (Platani et al., 2018; Platani et al., 2009; Zuccolo et al., 2007). Whether these defects are caused by the mislocalization of the entire Y-complex from kinetochores, as observed in HeLa cells (Platani et al., 2009; Zuccolo et al., 2007) or by the removal of Seh1 alone has recently been questioned (Platani et al., 2018). To study the mitotic role of Seh1 in mESCs, *WT* and *Seh1^-/-^* cells were transfected with GFP-H2B and imaged for 4-6 hours. Quantification of progression time from prometaphase to anaphase onset showed a ∼10 min delay in *Seh1^-/-^* as compared to *WT* mESCs (from 23.7 ± 10.1 min in *WT* to 32.8 ± 14.5 min in Seh1*^-/-^* cells; mean±SD) (Fig. S2 A). This delay is clearly milder than the one initially reported upon RNAi-induced depletion of Seh1 in HeLa cells (∼45 to 60 min; Platani et al., 2009; Zuccolo et al., 2007) but comparable to the delay recently measured upon conditionally-induced degradation of Seh1 in a HCT116-derived cell line (∼12 min; Platani et al., 2018). In *Seh1^-/-^* or auxin-treated *GFP-mAID-Seh1* mitotic mESCs, the Y-complex (visualized by Nup133 and Nup85) was still properly localized at kinetochores despite the complete lack of Seh1 (**Figs. S1C** and **S2 B,C**). This indicates that the mitotic delay observed in Seh1-deficient mESCs is not merely caused by the mislocalization of the Y-complex from kinetochores.

The 10-minute prolongation of mitosis was however unlikely to explain the cell growth defect of *Seh1^-/-^* mESCs (**Fig. 1C,D**). We therefore also measured the length of interphase by imaging mCherry-H2B-expressing mESCs during 24-30 hours. Quantification of progression time from the end of one mitosis (set at anaphase onset) to the beginning of the next (set at prometaphase) showed that interphase length is significantly longer in *Seh1^-/-^* as compared to *WT* mESCs (9.4 ± 2.2 hours in *WT* versus 14.0 ± 4.5 and 14.1 ± 2.1 hours in *Seh1^-/-^* #1 and #2, respectively; means ± SD) (**Fig. 2A**).

**Figure 2:**
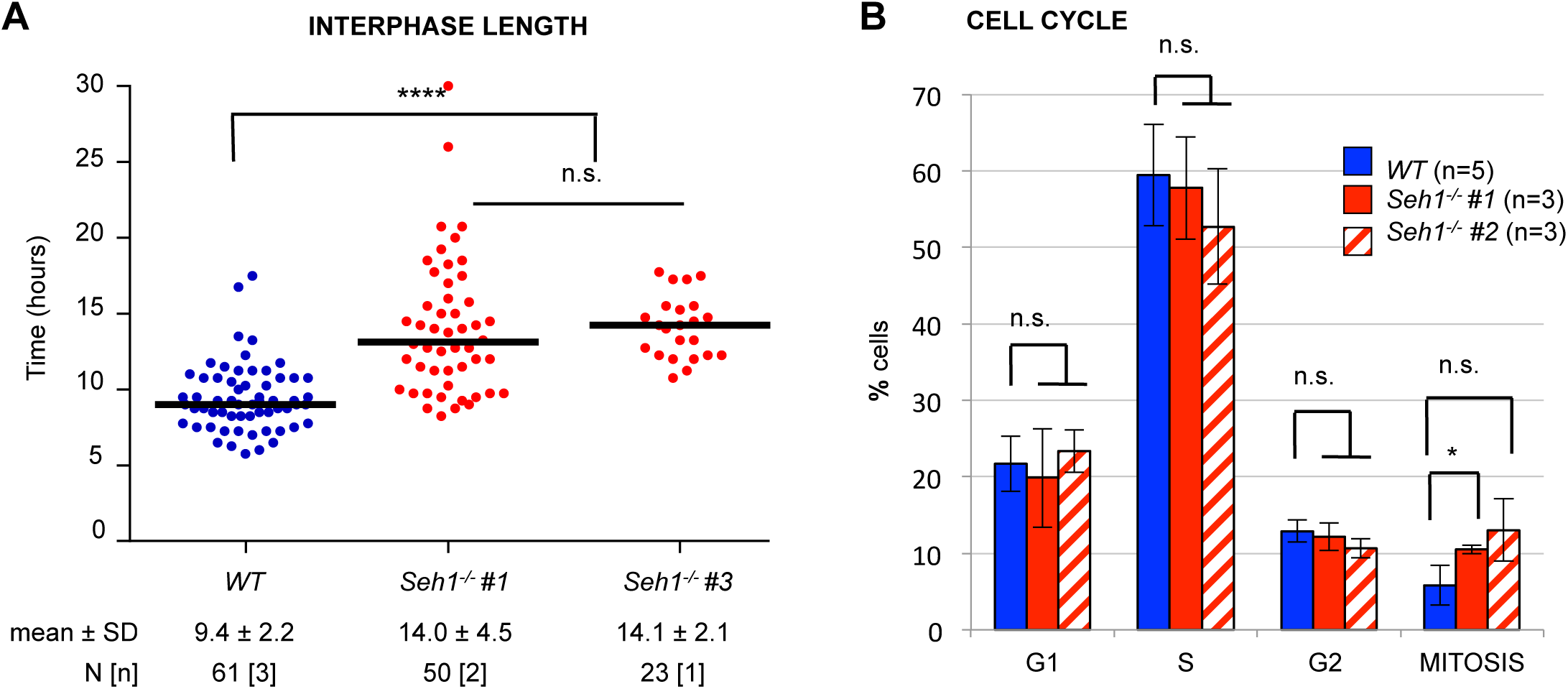
The altered growth rate of *Seh1^-/-^* mESCs reflects a lengthening distributed over all phases of the cell cycle. **A.** Quantification of interphase length of *WT* and *Seh1^-/-^* mESCs (2 distinct clones). The black bars represent the median and each dot represent one individual cell. The mean duration of interphase, as well as the number of imaged cells (N) and experiments [n] is indicated **B.** Cell cycle profiles of *WT* and two *Seh1 ^-/-^* clones generated by ImageStream→ using the workflow analysis presented in **Fig. S2D**. For each cell line, at least 3000 cells acquired in at least 3 distinct experiments were analyzed.

To determine if the lengthening of interphase in *Seh1^-/-^* mESCs was caused by retention in a specific phase of the cell cycle, we analyzed EdU-labelled and DAPI-stained *WT* and *Seh1^-/-^* mESCs by imaging flow cytometry (**Figs. 2B** and **S2D**). Except for a mild increase in the percentage of the mitotic fraction, this analysis revealed a comparable distribution of the G1, S and G2 phases of the cell cycle between *Seh1^-/-^* and *WT* mESCs (**Fig. 1H**). Therefore, the altered growth rate of *Seh1^-/-^* mESCs reflects a lengthening distributed over all phases of the cell cycle.

### Lack of Seh1 leads to a decrease of both NPC density and nuclear size

The viability of *Seh1^-/-^* mESCs at the pluripotent stage and their impaired survival upon differentiation was reminiscent of the phenotype observed upon inactivation of Nup133, another member of the Y-complex (Lupu et al., 2008). Because Nup133 loss was recently demonstrated to affect NPC basket assembly (as revealed by lack of TPR staining in about 50% of the NPCs) (Souquet et al., 2018), we decided to examine the impact of *Seh1* inactivation on NPC assembly. We therefore quantified the average fluorescence intensity at the nuclear envelope (NE) of TPR, Nup98 and Nup133 in *WT* and *Seh1^-/-^* mESCs, using a GFP-tagged cell line for internal reference, as previously reported (Souquet et al., 2018; see Materials and Methods). This analysis revealed a mild but significant reduction (in the range of 20-35%) of the signal of these three nucleoporins in *Seh1^-/-^* relative to *WT* mESCs (**Fig. 3 A-C**). The fact that the reduced intensity at the NE is not restricted to TPR indicates that, unlike what was previously observed in *Nup133^-/-^* mESCs, the lack of Seh1 leads to a decrease in the total number of NPCs rather than alteration of a specific substructure. This defect in NPC density was also observed upon auxin-induced depletion of Seh1 (**Fig. 3D**) and was largely rescued by stable expression of GFP-Seh1 (**Fig. 3, A-C**).

**Figure 3:**
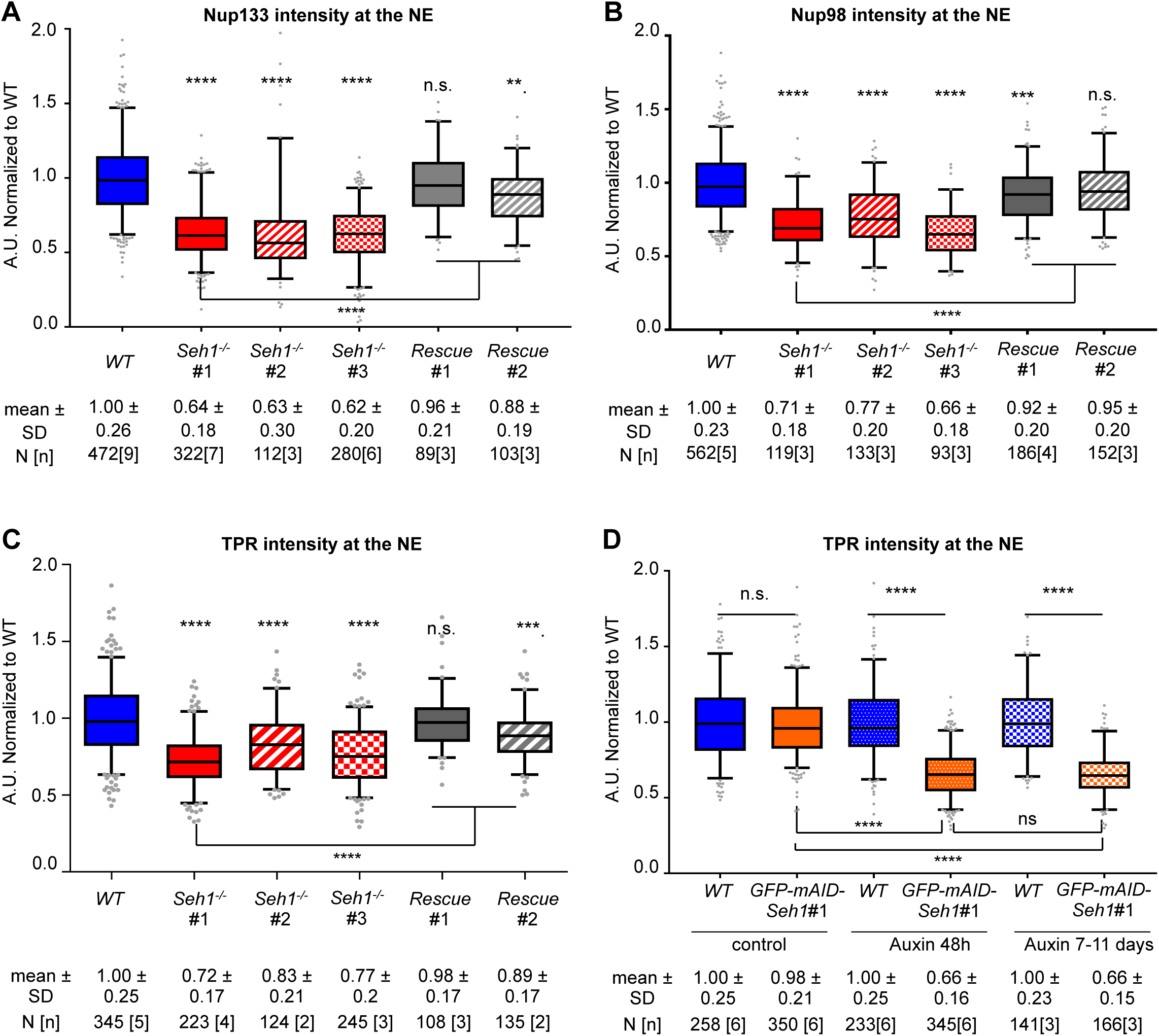
Quantification of NPC density in *Seh1* mutant cell lines. Normalized signal intensities at the NE of Nup133 (**A**), Nup98 (**B**) and TPR (**C-D**) were quantified and box plots generated as described in Materials and Methods. The mean value was set at 1 for *WT* mESCs. For each cell line, the number of cells quantified (N), the number of distinct experiments [n], the mean value and standard deviation are indicated. In (**D**), cells were treated with EtOH (control) or Auxin as indicated.

It was recently proposed that nuclear size is sensitive to NPC density and nuclear import capacity in cultured mammalian cells (Jevtić et al., 2019). The decreased NPC density observed upon Seh1 inactivation thus prompted us to measure nuclear size in these mutant mESCs. This analysis revealed a ∼10% reduction of the nuclear surface in *Seh1^-/-^* mESCs, a phenotype that was rescued by the GFP-Seh1 transgene (**Fig. 4A**). A significant reduction in nuclear size could also be observed in auxin-treated *GFP-mAID-Seh1* mESCs (**Fig. 4B**).

**Figure 4:**
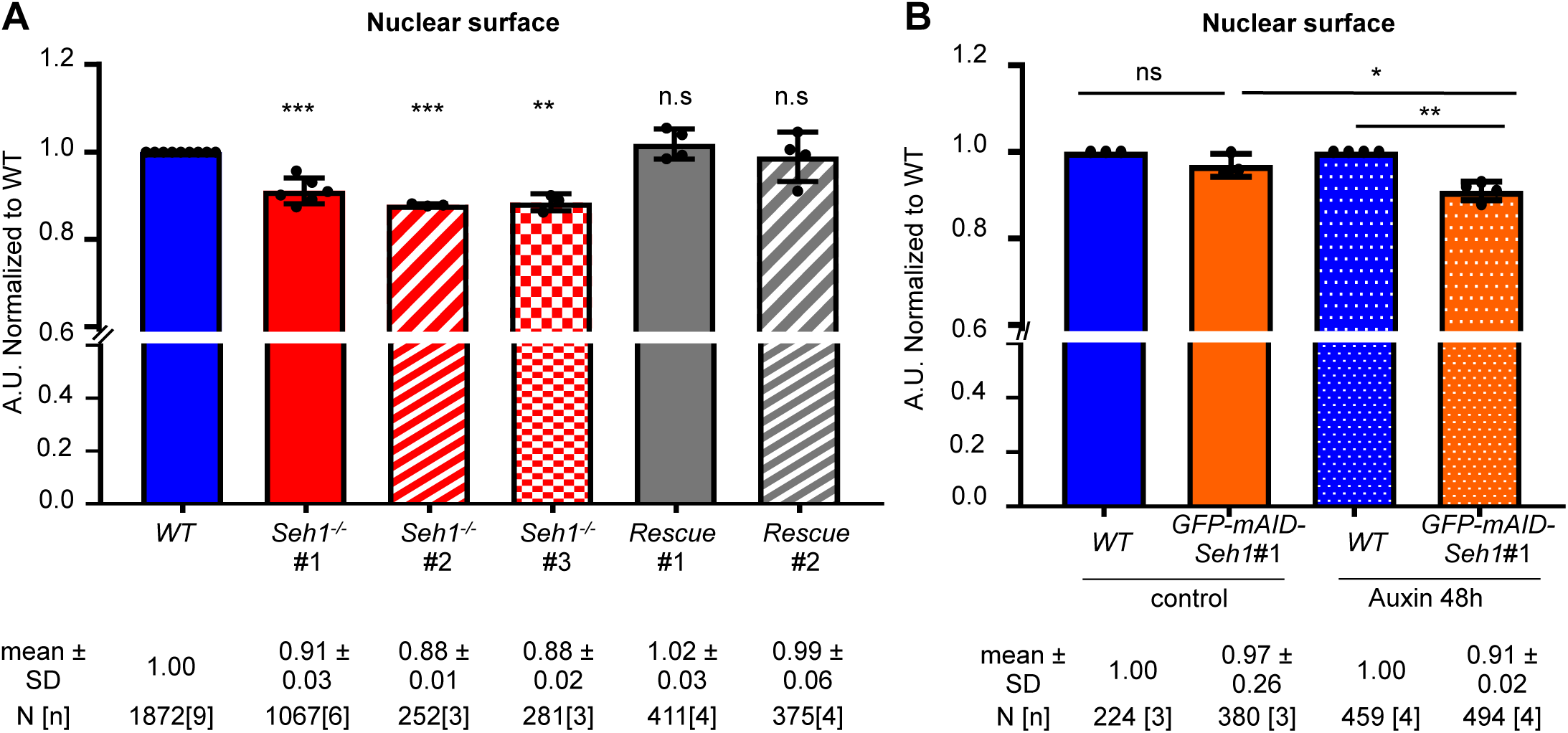
Quantification of nuclear sizes in *Seh1* mutant cell lines. Quantification of nuclear surface was performed as described in Materials and Methods. The mean value was set at 1 for *WT* mESCs. Graphs show average and standard deviation of nuclear surface values from [n] independent experiments (displayed as dots). Unless specified by lines, samples were compared to *WT* for statistical analyses (see Materials and Methods)

*Seh1*-deficient mESCs thus exhibit several distinct phenotypes: altered cell growth, lethality upon differentiation, reduced NPC density and nuclear size. We next aimed to determine whether these defects were linked to each other and whether they reflected functions of Seh1 as part of the Y-complex or the GATOR2 complex, or both.

### Mios is not required for proper cell growth and cell differentiation in mESCs

Within the GATOR2 complex, Seh1 directly interacts with Mios (also known as Mio, <u>mi</u>ssing <u>o</u>ocyte in Drosophila and Sea4 in budding yeast) (Senger et al., 2011; Bar-Peled et al., 2013; Algret et al., 2014). Our western blot analyses revealed decreased protein levels of Mios in *Seh1^-/-^* compared to *WT* mESCs (**Fig. 5A)**, a result consistent with studies in other species and cell types (Platani et al., 2018; Platani et al., 2015; Senger et al., 2011). To assess if this reduction in Mios could cause the cell growth and differentiation phenotypes observed in *Seh1^-/-^* mESCs, we inactivated *Mios* in mESCs via CRISPR/Cas9 (**Fig. 5A** and **Table S2**). Immunoprecipitation experiments performed using anti-Seh1 antibodies revealed that lack of Mios prevents Seh1 interaction with Wdr24, another GATOR2 complex component (Bar-Peled et al., 2013) (**Fig. 5B**). This points to Mios as being the main direct partner linking Seh1 to the rest of the GATOR2 complex, a result complementing data previously obtained in budding yeast and drosophila (Algret et al., 2014; Dokudovskaya and Rout, 2015; Cai et al., 2016). Analyses of independent *Mios^-/-^* clones revealed only a minor reduction of cell growth at the pluripotent stage as compared to *WT* mESCs (14±15% decrease in confluence after 48h of growth, while the reduction was 44±14% for *Seh1^-/-^* mESCs; **Fig. 5C**). In addition, *Mios^-/-^* cells underwent differentiation towards the neuroectodermal lineage with a comparable cell density (**Fig. 5D**) and morphology (our unpublished data) as *WT* cells. Finally, quantitative analyses did not reveal any significant alteration in either NPC density or nuclear size in *Mios^-/-^* as compared to *WT* mESCs (**Fig. 5E, F**).

**Figure 5:**
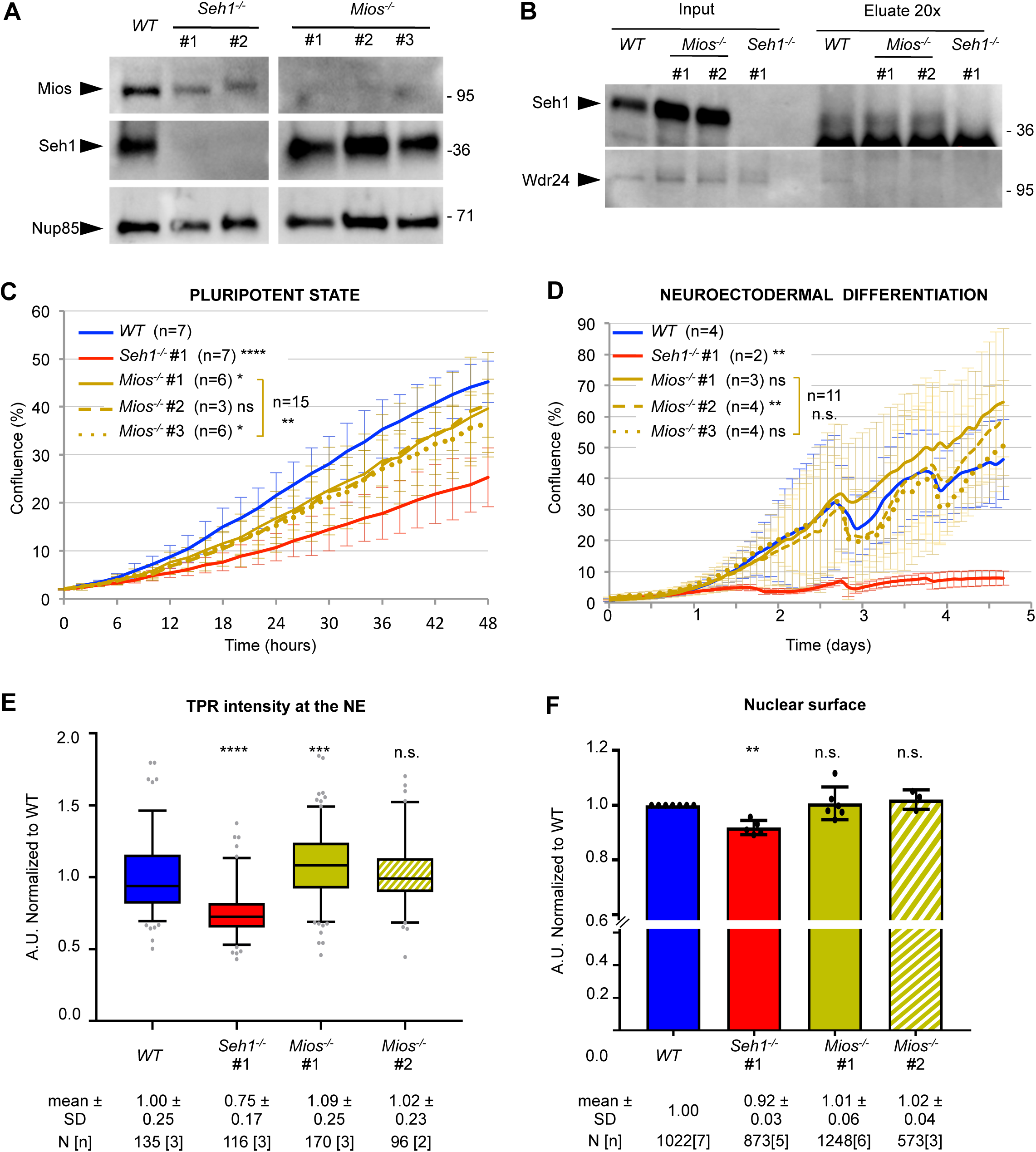
*Mios^-/-^* mESCs do not mimic the growth and differentiation defects of *Seh1^-/-^* mESCs, nor their decreased NPC density and nuclear size. **A.** Whole cell extracts of *WT*, *Seh1^-/-^* (#1 and #2) and *Mios^-/-^* mESCs (three independent clones) were analyzed by western-blot using the indicated antibodies. Molecular masses are indicated on the right (kDa). **B**. Immunoprecipitation experiment using anti-Seh1 antibodies and *WT*, *Mios^-/-^* (#1 and #2), or *Seh1^-/-^* (#1) mESC protein extracts. Inputs and eluates (20x equivalent) were analyzed by western blot using the indicated antibodies. Molecular markers are indicated on the right (kDa). **C-D.** Cell growth analyses (using percentage of confluence as proxy) were performed with the IncuCyte→ system for *WT*, *Seh1^-/-^* and three distinct *Mios^-/-^* clones at pluripotent state (**C**), and upon neuroectodermal differentiation (**D**). Error bars correspond to the standard deviation arising from the indicated number of independent experiments [n]. Statistical analyses were performed at the last time-points. Brackets indicate statistics performed using all values from cell lines bearing a given mutation, compared to *WT* (see Materials and Methods). **E-F.** Quantifications of TPR signal intensity at the NE (**E**, presented as box plots) and of the nuclear surface (**F**, graphs presenting the mean values and standard deviations from [n] distinct experiments, each displayed as a dot) were performed for *WT*, *Seh1^-/-^* and two *Mios^-/-^* clones as described in Materials and Methods. For each cell line, the total number of cells (N) acquired in [n] distinct experiments, and the mean and standard deviation values are indicated. For statistical analyses (see Materials and Methods) samples were compared to *WT.* Note that the mild (9%) <u>increase</u> in TPR density in *Mios^-/-^* # 1 mESCs was not observed for *Mios^-/-^* #2 cells and likely reflects a clonal-related variation not linked to the lack of Mios. Note that data for *WT* and *Seh1^-/-^* mESCs (used as reference strains) shown in panels C, D and F, include some data from experiments already presented in Figs. 1D-E and 4A.

Together these experiments indicate that neither the growth and differentiation defects, nor the altered nuclear pore density and nuclear sizes observed in *Seh1^-/-^* mESCs can be merely attributed to the decreased levels of Mios.

### Mutations affecting the short arm of the Y-complex impair NPC assembly, but with distinct impacts on cell proliferation and differentiation

Having excluded Mios destabilization as the main cause of the defects of *Seh1^-/-^* mESCs, we next focused our attention on Seh1’s partners localized on the short arm of the Y-complex (**Figs. 1A** and **6A**).

We first inactivated *Nup43*, another small β-propeller-folded nucleoporin that is specific to metazoan Y-complexes (Neumann et al., 2010). We obtained viable clones upon CRISPR/Cas9-mediated *Nup43* knockout in mESCs that however displayed impaired proliferation at the pluripotent stage (**Fig. 6B, C**). Although this growth defect was milder than the one observed in pluripotent *Seh1^-/-^* mESCs, *Nup43^-/-^* nevertheless underwent drastic cell death upon neuroectodermal differentiation, comparable to that of differentiating *Seh1^-/-^* cells (**Fig. 6D**). *Nup43^-/-^* mESCs also displayed a reduced NPC density comparable to that observed in the various *Seh1^-/-^* mESC lines, as revealed by the reduced intensity of Nup133, Nup98 and TPR labelling at nuclear pores (**Fig. 7A-C**, see also **Fig. 2A-C**). Finally, these cells showed no significant reduction in nuclear size (**Fig. 7D**). Together these data indicate that inactivation of *Nup43* mimics, albeit with a slightly milder impact, most of the phenotypes caused by *Seh1* inactivation.

**Figure 6:**
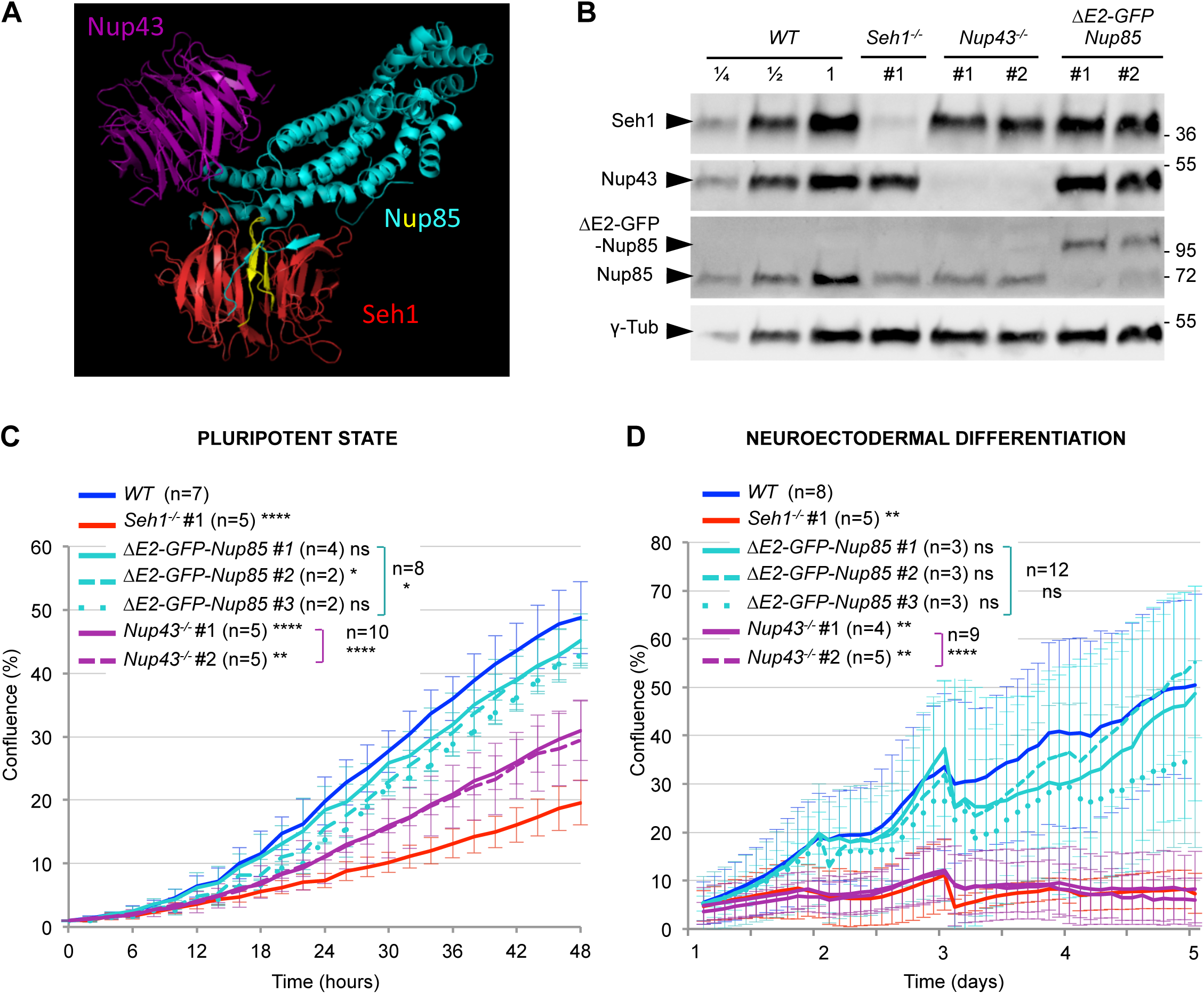
Impact of Y-complex short arm mutations on mESC proliferation and differentiation. **A.** Predicted model of human Nup43 (shown in purple), Nup85 (blue and yellow) and Seh1 (red) interactions (von Appen et al., 2015; PDB code: 5A9Q) visualized using Pymol. The β-sheets within the N-terminal domain of Nup85 that are deleted in the ΔE2-GFP/mCherry-Nup85 fusions are shown in yellow. **B.** Whole cell extracts of the indicated cell lines were analyzed by western-blot using anti-Seh1, -Nup43, -Nup85, and γ-tubulin antibodies. Two-and four fold dilution (1/2, 1/4) of the *WT* mESC extract were also loaded. Molecular markers are indicated on the right (kDa). **C-D.** Cell growth analyses were performed with the IncuCyte→ system for *WT*, *Seh1^-/-^*, *ΔE2-GFP-Nup85* and *Nup43^-/-^* mESCs at pluripotent state (**C**), and upon neuroectodermal differentiation (**D**). Error bars correspond to the standard deviation arising from [n] independent experiments. Statistical analyses were performed at the last time-points. Brackets indicate statistics performed using all values from cell lines bearing a given mutation, compared to *WT* (see Materials and Methods). Note that data for *WT* and *Seh1^-/-^* (used as reference strains) shown in panels **C-D** include experiments already presented in Fig. 1 D-E.

**Figure 7:**
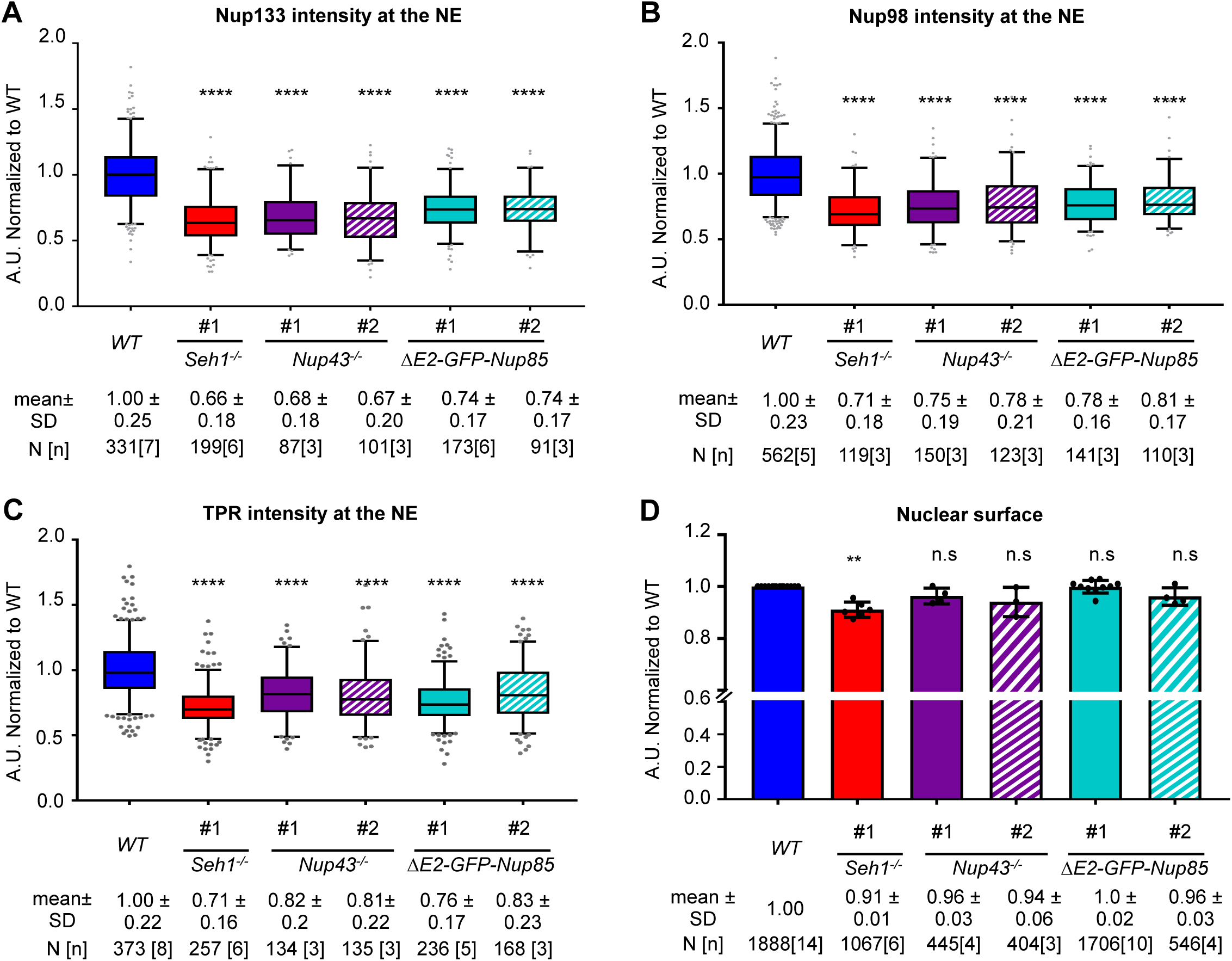
Quantification of NPC density and nuclear size in *ΔE2-GFP-Nup85* and *Nup43^-/-^* mESC lines. **A-C.** Normalized signals intensities at the NE of Nup133 (**A**), Nup98 (**B**) and TPR (**C),** (presented as box plots) and nuclear surfaces (**D;** graphs presenting the mean values and standard deviations from [n] distinct experiments, each displayed as a dot) were quantified as described in Materials and Methods. The number of cells (N), and of distinct experiments [n], the mean value and standard deviation are indicated. For statistical analyses (see Materials and Methods) samples were compared to *WT.* Note that data for *WT* and *Seh1^-/-^* (used as reference strains) include experiments already presented in Fig. 3.

To determine whether these shared phenotypes reflect a function of these Nups within the short arm of the Y-complex, we next aimed to impair Seh1 recruitment to the NPCs. Structural studies have shown that budding yeast Seh1 binds to Nup85 through its N-terminal domain forming the 7^th^ blade to complete the β-propeller structure of Seh1 (Brohawn et al., 2008; Debler et al., 2008). Homology modelling predicted that the human Nup85-Seh1 interface similarly involves ß-sheets within Nup85 N-terminal domain that complete Seh1 ß-propeller (von Appen et al., 2015); see **Fig. 6A**). We thus attempted to prevent Seh1 binding to Nup85 by deleting most of this blade (two β-sheets encoded by exon 2 of mouse Nup85; colored in yellow in **Fig. 6A**) and inserting instead the sequence of the bulky GFP. We obtained viable mESC lines in which the resulting ΔE2-GFP-Nup85 fusion, expressed as the unique form of Nup85 in the cell (**Figs. 6B** and **S3C)**, was properly localized at both NPCs and kinetochores (**Fig. S3A**).

To determine whether this deletion within Nup85 indeed prevented its interaction with Seh1, we performed immunoprecipitation on *WT* or *ΔE2-GFP-Nup85* mESC lysates using antibodies directed against either Nup85 itself, or Nup107, another Y-complex constituent (**Fig. 1A**). Mass-spectrometry analysis showed that ΔE2-GFP-Nup85 interacts with all the members of the Y-complex except Seh1 (**Fig. S3B**). Because none of the available Seh1 antibodies we tested properly recognized endogenous mouse Seh1 in immunofluorescence experiments, we also introduced within *GFP-Seh1* cells the same N-terminal deletion of Nup85, this time tagged with mCherry (**Fig. S3C** and **Table S2**). Although ΔE2-mCherry-Nup85 was properly localized at NPCs and kinetochores (**Fig. 8A,B**), GFP-Seh1 was at most only barely detectable at kinetochores in these cells (**Fig. 8B,D**; quantifications revealed 7-8 ± 5-7% residual signal in ΔE2-mCherry-Nup85-compared to wt-Nup85-expressing cells). The mislocalization of Seh1 from kinetochores is consistent with its impaired interaction with ΔE2-GFP-Nup85 seen at the biochemical level and with previous studies indicating that the Y-complex is recruited as an entity to kinetochores (Loiodice et al., 2004). In contrast, we could still detect some punctate GFP-Seh1 signal at the nuclear envelope (26-27% residual signal; **Fig. 8A,C**). The relative persistence of GFP-Seh1 at NPCs as compared to kinetochores in *ΔE2-mCherry-Nup85* cells likely reflects the existence of additional minor binding sites for Seh1 at NPCs, that either involve Nups not belonging to the Y-complex, or imply interfaces generated by the 3D-organization of Y-complex within the assembled mammalian NPC (Huang et al., 2020; Kosinski et al., 2016; von Appen et al., 2015).

**Figure 8.**
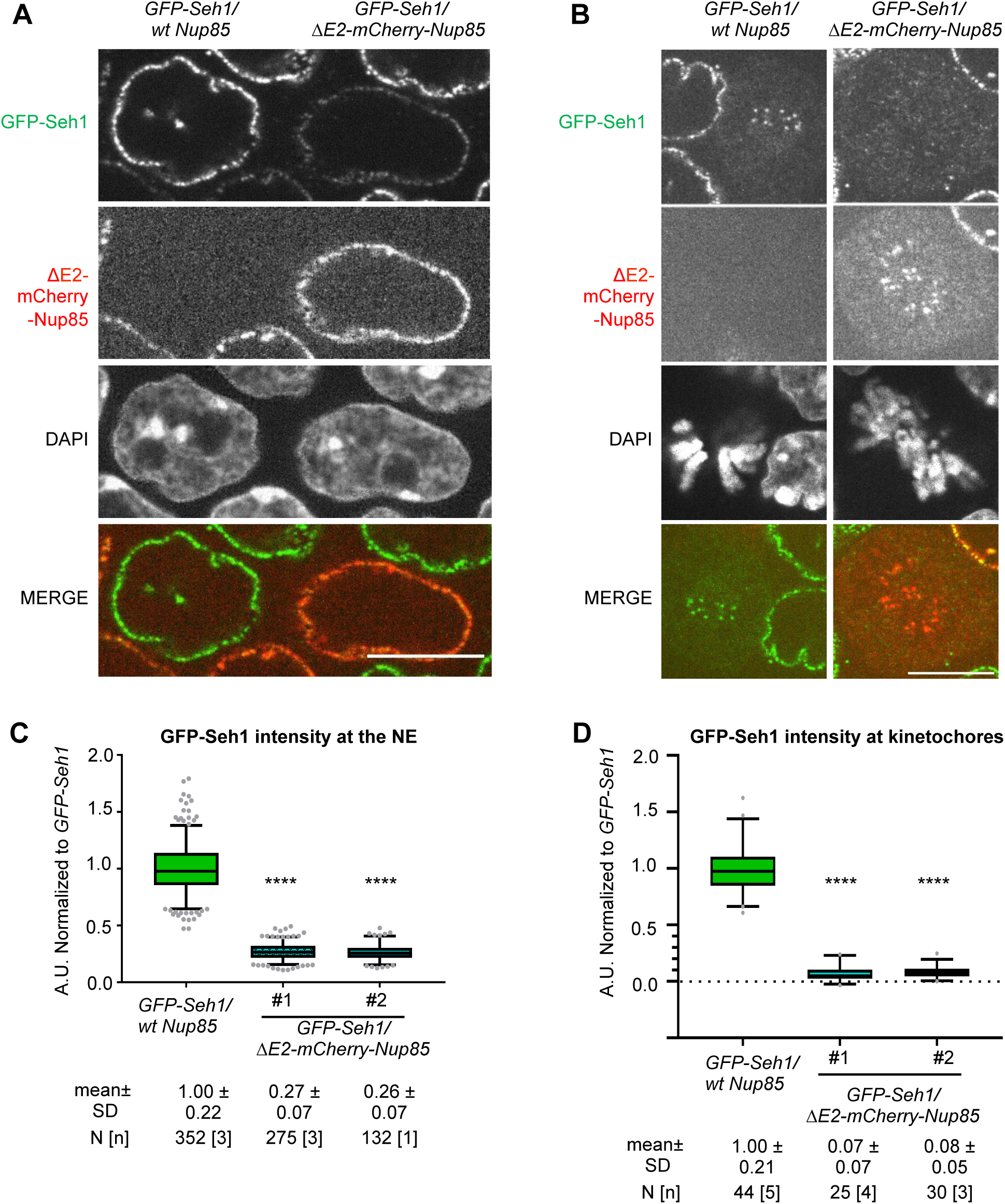
Impaired GFP-Seh1 localization at NPCs and Kinetochores in *ΔE2-mCherry-Nup85* mESCs. **A-B.** Representative spinning disk images (single z-section) of interphase (**A**) and mitotic (**B**) *GFP-Seh1* cells (left) mixed with *ΔE2-mCherry-Nup85/GFP-Seh1* cells (right). Scale bars, 10 μm. **C-D.** GFP-Seh1 intensity at NE (**C**) and at kinetochores (**D**) was quantified in *GFP-Seh1* and *ΔE2-mCherry-Nup85/GFP-Seh1* for the indicated number of cells (N) acquired in [n] independent experiments as described in Materials and Methods. Values were normalized for each field (**C**) or for each experiment (**D**) to the average intensity of the signal acquired for *GFP-Seh1* cells at the NE and kinetochores, respectively.

Unexpectedly, analysis of cell differentiation did not reveal any significant differences between *ΔE2-GFP-Nup85* and *WT* mESCs (**Fig. 6D**). In addition *ΔE2-GFP-Nup85* mESCs only displayed a minor cell growth defect in the pluripotent state (9±7% decrease in confluence after 48h of growth compared to *WT* mESCs, while the reduction was 60±7% and 38±11% for *Seh1^-/-^* and *Nup43^-/-^* cells, respectively; **Fig. 6C**). Nevertheless, pluripotent *ΔE2-GFP-Nup85* cells display a significant reduction in the intensity of Nup133, Nup98 and TPR at the nuclear envelope (**Fig. 7A-C**). These decreased signals were comparable to those observed in *Seh1^-/-^* and *Nup43^-/-^* mESCs, and yet, as in the case of *Nup43^-/-^*, they were not accompanied by a significant reduction in nuclear size (**Fig. 7D**). Analysis of the *ΔE2-GFP-Nup85* cells lines thus showed that perturbed recruitment of Seh1 at NPCs leads to a reduction in NPC number, but does not impact cell growth and differentiation.

## Discussion

This study has revealed that Seh1 and Nup43 are dispensable for mESC viability in the pluripotent state but become critical upon their differentiation. In view of the reported embryonic lethality of the *Seh1* knockout in mouse, an impaired differentiation of *Seh1^-/-^* mESCs could have been anticipated (Liu et al., 2019). In contrast, no role in development had been previously described for Nup43, which is specific to metazoans (Neumann et al., 2010). Although the requirement for differentiation was reminiscent of the phenotype observed upon inactivation of *Nup133,* we observed that Seh1 and Nup43, are, unlike Nup133 (Lupu et al., 2008), also required for proper growth of mESCs in the pluripotent state. Importantly, this altered growth is not simply caused by a mitotic defect, as might have been assumed given the mitotic roles of Seh1 in cancer cells (Platani et al., 2018; Platani et al., 2009; Zuccolo et al., 2007), but rather reflects a lengthening of all phases of the cell cycle.

We initially anticipated that the phenotypes of *Seh1^-/-^* mESCs could be caused by a combination of its functions within the Y- and the GATOR2-complexes. However, the fact that the *Mios^-/-^* cells do not feature any NPC assembly, nuclear size, or cell differentiation defects rather suggests that *Seh1^-/-^* phenotypes (except perhaps a mild contribution to cell growth rates) are unlikely to result from a combination of defects in NPC and GATOR function. Moreover, *Nup43^-/-^* mESC phenotypes are very similar to those of *Seh1^-/-^* mESCs, despite the fact that Nup43 does not interact with Mios.

Our data also showed that integrity of the short arm of the Y-complex is important for proper NPC density, further distinguishing its function from that of Nup133, which specifically affects NPC basket assembly in mESCs (Souquet et al., 2018). The observed reduction in NPC density in our *Seh1*^-/-^, *Nup43^-/-^* and *ΔE2-GFP-Nup85* clones likely reflects an absolute reduction in total NPC number, since there was no corresponding increase in nuclear surface in these cells (instead, nuclear size was mildly reduced in *Seh1^-/-^* mESCs). Different mechanisms may explain the requirement for an intact short arm of the Y-complex to ensure proper NPC numbers. Considering the critical roles of the Y-complex in both NPC re-assembly after mitosis and *de novo* NPC assembly in interphase (Doucet et al., 2010; Harel et al., 2003; Walther et al., 2003), the short arm of the Y-complex might be required for the efficient recruitment of the Y-complex either to the mitotic chromatin (an hypothesis consistent with the minor reduction of Y-complex levels on chromatin reported upon Seh1 depletion in HCT116 cells- Platani et al., 2018), or to the nuclear envelope in interphase (as process involving Nup153 Vollmer et al., 2015). Alternatively, Nup43 and Seh1 may contribute to the stabilization of the NPC scaffold, by virtue of their direct interactions with neighbouring subunits from either Y-complexes or inner ring complexes (Huang et al., 2020; Kosinski et al., 2016; von Appen et al., 2015). NPCs lacking these stabilizing interactions might then be recognized by one of the recently described quality-control mechanisms that mediate the removal of some misassembled NPCs from the nuclear envelope (reviewed in Webster and Lusk, 2016).

Finally, our analysis of the *ΔE2-GFP-Nup85* cell lines indicates that the reduction in NPC density observed in *Seh1*^-/-^ and *Nup43^-/-^* mESCs is not sufficient to impact cell growth and differentiation. The lack of major growth and differentiation defects in *ΔE2-GFP-Nup85* cells, in which Seh1 is largely mislocalized from NPCs, could reflect an “off-pore” function of Seh1, or a function of Seh1 that does not require its normal stoichiometry within NPCs (for instance, a localization restricted to the cytoplasmic or nuclear side of the NPCs). At NPCs, Seh1 and Nup43 might be required for the proper recruitment and positioning of the mRNA export and remodelling machinery, an established function of the short arm of the Y-complex in budding yeast (Fernandez-Martinez et al., 2016). Alternatively, whether at pores or elsewhere in the nucleus, Seh1 and Nup43 may impact cell growth and differentiation by directly contributing to gene regulation, as now reported for a few Nups in mammalian cells (reviewed in Buchwalter et al., 2019; Pascual-Garcia and Capelson, 2019; see also Scholz et al., 2019). In particular, Seh1 was recently found to participate in oligodendrocyte differentiation, acting as a platform to recruit transcription and chromatin remodelling factors (Olig2 and Brd7) (Liu et al., 2019). We may hypothesize that both Seh1 and Nup43 may specifically interact with factors required for gene regulation and chromatin organization in mESCs, hence contributing to early stages of pluripotent cell growth and differentiation.

## Materials and Methods

**Plasmids** used in this study are listed in **Table S1**. They were either previously published or generated using standard molecular cloning techniques including restriction digestions (Fastdigest, Thermo Fisher Scientific, Waltham, MA), PCR amplification using proofreading DNA polymerases (Phusion HF, New England Biolab, Ipswich, MA) and In-Fusion HD Cloning Kit (Clontech, Mountain View, CA) or NEBuilder HiFi DNA Assembly Cloning Kits (New England Biolab). The *Mios* and *Nup43* gRNAs were integrated in a linear plasmid (GeneArt™ CRISPR Nuclease Vector – OFP-Cas9) following manufactureŕs instructions. The other Cas9 vectors (pX-280, pX-672, pX-853 and pX-864) were assembled by golden gate cloning (Engler et al., 2009). For all constructs, PCR-amplified fragments and junctions were checked by sequencing. Plasmid maps are available upon request.

### Cell lines, growth condition, transfection, and CRISPR/CAs9-based genome editing

Cell lines used in this study are listed in **Table S2**. All cells were grown at 37°C and 5% CO_2_.

<u>DR4-mouse embryonic fibroblast feeder cells</u>, DR4-MEFs (Applied StemCells), were grown in Dulbecco’s Modified Eagle’s Medium (DMEM) (Gibco/Thermofisher) supplemented with 15% heat-inactivated foetal bovine serum (FBS, Gibco), 100U/ml Penicillin-100µg/ml Streptomycin (P/S) (Gibco) and 2mM L-Glutamine (Gibco). DR4-MEFs were inactivated using 8.5 µg/ml Mitomycin-C (Sigma-Aldrich, St Louis, MO) for 3 hours.

<u>HM1</u> (Selfridge et al., 1992) <u>and derivative mESCs clones</u> were grown in serum/leukemia inhibitory factor (LIF)-containing stem cell medium: DMEM (EmbryoMax, Millipore, Burlington, MA), P/S (Gibco), 2mM L-Glutamine (Gibco), 15% heat-inactivated ESC-Qualified FBS (Gibco), non-essential amino acids (Gibco), nucleosides (Millipore), 2-mercaptoethanol (Gibco) and 10^3^ units/ml LIF (ESGRO, Millipore). mESCs were grown on inactivated DR4-MEFs (MEF-derived feeders) plated on 0.1% gelatin (Sigma-Aldrich) and were passaged every 2 or 3 days using 0.05% Trypsin (Gibco). mESCs were used at passages below 30. Lack of contamination in-between the mutant cell lines was assessed by PCR on genomic DNA, proper GFP-or mCherry expression when pertinent, and western blots analyses. Frequent DAPI staining ensured lack of major contamination by mycoplasm. When required, cells were counted using a Countess automated cell counter (Invitrogen, Carlsbad, CA).

<u>For transfections</u>, mESCs were plated onto DR4-MEFs in medium without P/S. Plasmid DNA and Lipofectamine 2000 (Invitrogen) were mixed in OptiMEM (Invitrogen) and added to the cells according to the manufacturer’s instructions.

<u>For CRISPR/Cas9 editing</u>, 5·10^5^ mESCs were transfected as indicated in **Table S2** with one or two plasmids (3µg each) directing the expression of one or two gRNAs along with Cas9 (WT or high fidelity, HF) fused to GFP, mCherry or OFP. gRNAs were designed using the Benchling website (https://benchling.com) and are listed in **Table S3**. When indicated, DNA sequences of interest (PCR product 1-4 µg, or 3µg of linearized plasmid) flanked by homology directed repair arms were co-transfected **(Fig. S4)**. Following selection (as detailed below), individual clones were picked, amplified, and further characterized. For each clone, chromosome spreads were also performed (chromosome counts are indicated in **Table S2**).

<u>To establish *Seh1^-/-^, Mios^-/-^* and *Nup43^-/-^* cell lines,</u> cells were collected by trypsinization two days after transfection, resuspended in 1 mL Fluorescence-activated cell sorting (FACS) buffer (PBS +10% FBS, Gibco + P/S), and sorted based on Cas9 expression (EGFP or OFP signal). 2000 FACS-sorted cells were plated in 100mm culture dishes. Individual clones picked 6-12 days after sorting were then characterized using Western blot and PCR on genomic DNA followed by sequencing (the identified Indels are listed in **Table S2**).

To establish the *<u>OsTir</u>* <u>cell line</u>, 200µg/mL Geneticin (Geneticin® Gibco, Life technologies 10131-019) was added to the medium two days after transfection. Geneticin-resistant clones (expected to have integrated the pCAG-OsTir-T2A-NeoR sequence at the *Tigre* locus) were picked after five days and characterized by Western-blot with antibodies directed against the OsTir receptor. PCR on genomic DNA was also performed to determine the number of *Tigre* alleles bearing the transgene.

To generate the *<u>Seh1 rescue</u>* (expressing GFP-Seh1 under the pCAG promoter at the *Tigre* locus), *<u>GFP-Seh1</u>*, *<u>GFP-mAID-Seh1</u>*, *<u>ΔE2-GFP-Nup85</u>* and <u>[*ΔE2-mCherry-Nup85*]</u> cell lines, GFP [mCherry] positive cells were FACS-sorted 3 days after transfection to select for cells expressing the tagged nucleoporin. Individual clones were picked 6-7 days after sorting and characterized using immunofluorescence (to confirm the localization of the tagged protein at the nuclear periphery) and western blot (to identify clones lacking the endogenous protein). The selected clones were then further validated by PCR on genomic DNA and sequencing.

To achieve inducible degradation of GFP-mAID-Seh1, Auxin (Sigma-Aldrich) was added to the medium at 500µM (from a stock at 280 mM in EtOH). For control experiments, the same amount of EtOH was added.

### Cell growth and differentiation assays

To evaluate <u>cell growth at pluripotent stage</u>, cells were plated at 1 −2·10^5^ cells per well in TPP 12-well plate. Photomicrographs were taken every two hours using an IncuCyte→ live cell imager (Essen Biosciences, Ann Arbor, MI) and confluence of the cultures was measured using IncuCyte→ software (Essen Biosciences, Ann Arbor, MI). To improve comparison in-between experiments or cell lines, the same mask was always used and time was set at t=0 when confluence reached 1% (**Figs. 1D and 6C**), 2% (**Fig. 5C**) or 3% (**Fig. S1E**). Graphs were generated using Excel. Error bars correspond to standard deviations from the indicated [n=] independent experiments.

<u>Neuroectodermal differentiation</u> of mESCs grown as monolayers was adapted from (Ying and Smith, 2003). Following trypsinization, feeders were removed by plating the resuspended cells in gelatin-free wells for 20 minutes. Feeder-free mESCs were collected and resuspended in N2B27 medium [DMEM F-12, DMEM Neurobasal, BSA, L-glutamin, 2-mercaptoethanol, N2 (Gibco) and B27 (Gibco)]. Cells were plated in gelatin-coated wells at 1·10^5^ or 3·10^4^ cells per well in TPP 12 well plate. At day 2, N2B27 medium containing 1μM RA (all-trans-Retinoic acid, Sigma) was added for 24 hours. From day 3 to 7 medium was changed every day with fresh N2B27 without RA. Confluence analyses, used as a proxy to evaluate cell growth and viability, was performed as described above except that time was set at t=0 the beginning of the differentiation process (i.e., upon plating in N2B27 medium).

### Fluorescence Videomicroscopy

mESCs were transiently transfected using plasmids expressing H2B-GFP or H2B-mCherry on microscopy-adapted 35-mm dishes (µ-dish, 35 mm, high; Ibidi, Germany) coated with 0.1% gelatin and DR4-MEFs. Acquisitions were performed about 36 hours after transfection at 37°C and 5% of CO_2_ using an AxioObserverZ1 inverted microscope (Zeiss, Germany) equipped with a 63 oil objective, a CSU-X1 spinning-disk head (Yokogawa, Japan), and a sCMOSPRIME 95B (Photometrics) camera.

The whole setup was driven with MetaMorph software (Molecular Devices, Sunnyvale, CA). Eleven Z sections with a step of 1µm were acquired at intervals of 5 minutes for the mitotic progression experiments (4-6 hours) and of 15 minutes for the cell cycle length experiments (24-30 hours). Laser intensity was set between 10-20% power and acquisition time was 500ms. The raw data were processed using ImageJ software (National Institutes of Health, Bethesda, MD). Images stacks were processed as max projections. Cells were tracked manually setting prometaphase at the moment at which chromatin starts to be seen condensed and anaphase at the first time point at which chromosome segregation is observed.

### FACS analyses

To perform a bi-parametric analysis of cell cycle based on DNA content (DAPI) and DNA synthesys (EdU) we used the Click-it-EdU Imaging kits (Invitrogen). 0.5 10^6^ mESC were plated on MEF-derived feeders plated on 0.1% gelatin 2 days before the experiment. Cells were incubated with EdU 50 μm for 15 minutes, then collected by trypsinyzation and plated on gelatin dishes for 20 minutes to remove feeders.

Cells were then collected and centrifuged, washed in PBS and then fixed in PFA 3% for 15 minutes. Cells were then permeabilized with PBS+ 0.2% Triton-X100 and washed with PBS + 2% SVF. Click reaction (30 minutes) was performed following manufacturer’s protocol. DNA was stained for 20 minutes (DAPI 5 μM, RNaseA 0.1 mg/mL, 1% SVF in PBS), then samples were centrifuged and resuspended in 60μl PBS +2% SVF.

Sample acquisition was achieved with the ImageStream→ X (Amnis, Austin, TX) imaging flow cytometer and captured using the ISX INSPIRE™data acquisition software. Images of 5.000–20.000 cells were acquired at 40x magnification using the following channels: Ch1= 430-470 nm, BF (bright field); Ch6= 720-800nm, SS (side scatter); Ch7= 430-505 nm, DAPI; Ch9= 570-595, BF; Ch11= 660-720 nm, EdU-AF647. A compensation matrix was generated using fluorescence controls and applied to all samples. Analysis was then performed with the IDEAS software as follows: 1) definition of cells in focus, based on the gradient RMS; 2) definition of singlets, according to area and aspect ratio; 3) definition of cells using contrast and gradient RMS; 4) definition of nucleated cells using DNA content; 5) cell cycle phases were then identified using DAPI and EdU intensity; 6) mitotic cells were finally defined according to DAPI bright detail intensity and DAPI area threshold (**Fig. S2 D**).

### Immunostaining and quantitative image analyses

mESCs grown on coverslips were washed with PBS, then fixed using 3% para-formaldehyde (VWR, Radnor, PA) for 20 min and washed again with PBS. For all conditions, cells were then permeabilized in PBS + 0.2% Triton X-100, 0.02% SDS (Euromedex, Souffelweyersheim, FR), and 10 mg/mL BSA (Sigma). Antibody hybridizations and washes were also performed in this buffer. Primary and secondary antibodies (listed in **Table S4**) were incubated for 1 hr at room temperature. Cells were then incubated 5 min with 280nM DAPI (Sigma) in PBS and mounted with Vectashield (Vector, Maravai Life Sciences, San Diego, CA). Images were acquired using 100x/1.4 oil objectives on inverted and motorized microscopes, either a DMI8 (Leica), equipped with a CSU-W1 spinning-diskhead (Yokogawa, Japan) and 2 Orca-Flash 4 V2+ sCMOS cameras (Hamamatsu), or an Axio Observer.Z1 (Zeiss), equipped with CSU-X1 spinning-diskhead (Yokogawa, Japan) and 2 sCMOS PRIME 95 cameras (Photometrics).

<u>Quantification of NPC density</u> at the nuclear envelope (NE) was performed essentially as described (Souquet et al., 2018). Briefly, mESCs of interest were mixed with a GFP cell line of reference (*Rescue-Seh1* or *ΔE2-GFP-Nup85* cells, used for normalization) and grown on coverslips for 24h prior to fixation and immunostaining. For each acquired image, one z section was selected; 8-pixel-thick regions of interest (ROIs) were drawn freehand on the NE of both GFP-negative and -positive (reference) mESCs. Following subtraction of background, the signal intensity at the NE for each cell was normalized to the average NE intensity measured for the GFP-positive mESCs acquired in the same field. All values were then divided by the mean normalized intensity of *WT* mESCs acquired in the same experiment. Box plots were generated using GraphPad Software: each box encloses 50% of the normalized values obtained, centred on the median value. The bars extend from the 5^th^ to 95^th^ percentiles. Values falling outside of this range are displayed as individual points. <u>For kinetochore quantifications</u>, mixed *GFP-Seh1* and *ΔE2-mCherry-Nup85/GFP-Seh1* mESCs grown on coverslips were fixed, permeabilized, and stained with DAPI. Fields containing both *GFP-Seh1* and *ΔE2-mCherry-Nup85/GFP-Seh1* mitotic cells were selected. For each mitotic cell, the mean intensities of five distinct kinetochores (regions of 10-pixels in diameter) and of two “background” regions in the mitotic cytoplasm (40-pixels in diameter) were measured on a unique z-section. Following background subtraction, the average intensity of GFP-Seh1 at kinetochores in each mitotic cell was normalized to the intensity measured for the control (*wtNup85/ GFP-Seh1*) mitotic cells acquired in the same experiment. Box plots were generated as described above.

<u>To quantify nuclear surfaces</u>, mESCs of interest were mixed with a GFP cell line of reference and grown on coverslips for 24 hr prior to fixation and immunostaining as described above. For each field, 33 to 45 optical sections (0.5 μm apart) were acquired and nuclei were segmented based on TPR immunostaining with the Fiji plugin Lime-Seg (Machado et al., 2019). A circular ROI was drawn within the nucleus of each cell in the field and the LimeSeg Plugin “Sphere Seg advanced” was run with the following parameters: D0: 4; Zscale: 7.143; range in D0 units: 2; real xy pixel size: 0.07; F pressure: 0.025 for TPR-Cy3 staining and 0.019 for TPR-Cy5. Segmented structures for which the “free edges” values were above 0 (segmentation could not close the structure), and those for which the Euler characteristic was not comprised between −4/+4 (aberrant structures very far from a spherical shape) were discarded. For each cell, the segmentation perimeter and TPR staining along the z-axis were compared to further validate proper segmentation (less than 8% of the identified structures, frequently corresponding to the merge of two closely apposed nuclei, were manually discarded at that stage). Nuclear surfaces and volumes were then exported. To compensate for variability occurring during fixation or IF processing, nuclear surface values were first normalized to the average of the GFP-reference cells acquired within the same coverslip, and then to the mean of *WT* mESCs acquired in the same experiment. Nuclear surface graphs were generated using GraphPad Software: average and standard deviation (boxes and bars) of nuclear surface are displayed, along with values for each experiment (dots).

### Western blot analyses

To prepare whole cell lysates, mESCs were lysed in 2× Laemmli lysis buffer (150-mM Tris-HCL (pH 6.8), 5 % (wt/vol) SDS, 25 % (vol/vol) glycerol, and 0.01 % (wt/vol) bromophenol blue). Lysates were incubated for 3 min at 95 °C, clarified by sonication (Bioruptordiagenode: 4 cycles of 30 s on/off, high power), and denatured again for 3 min at 95°C. Protein concentration was then determined using a BCA assay kit (Thermo Fisher Scientific). Total protein extracts supplemented with β-mercaptoethanol (750 mM final, Sigma-Aldrich) were analysed by western blot. 10 µg of mESC lysate were separated on 4–12% or 10% SDS-PAGE gels (pre-cast NuPage® GE healthcare or Mini-Protean TGX Stain free precast gels, Biorad, Hercules, CA) and transferred to nitrocellulose (GE healthcare). The resulting blots were stained using Ponceau, saturated with TBS buffer + 0.1% Tween and 5% dried milk, and incubated in TBS + 0.1% Tween and 5% dried milk with primary antibodies, followed by either HRP-conjugated secondary antibodies of interest or HRP-conjugated anti-rabbit TrueBlot® secondary antibody (used in **Fig. 5B** to prevent interference from the denatured/reduced heavy and light chains of the anti-Seh1 antibody used for immunoprecipitation (primary and secondary antibodies are listed in **Table S4**). Signals were detected by enhanced chemiluminescence (SuperSignal® Pico or Femto, ThermoScientific) using ChemiDoc (Biorad).

### Immunoprecipitation experiments and mass spectrometry analyses

#### Immunoprecipitation experiments

Protein G beads (GE Helathcare) were washed three times with Wash buffer (100mM NaCl, 1mM EDTA, 25mM TRIS pH 7.5, 1mM DTT + protease inhibitor [Pi] solution); (bead centrifugations were performed at 500g 4°C). 30µL of beads were then incubated for 2 hours at 4°C in 250µL wash buffer containing 5µL of rabbit anti-Seh1 antibody (for **Fig. 5B**), or 25µL of rabbit polyclonal anti-Nup107 or anti-Nup85 serum or a pre-immune rabbit serum as control (for **Fig. S3B**). Antibodies used are listed in **Table S4**. After incubation, beads were washed 4 times with Wash buffer.

In the meantime, lysates were prepared from *WT, Seh1^-/-^*, *Mios^-/-^* or *ΔE2-GFP-Nup85* mESCs by resuspending frozen pellets of 4·10^6^ mESCs (∼500μg total proteins) in 200μl Lysis buffer (100mM NaCl, 1mM EDTA, 25mM TRIS pH 7.5, 1mM DTT, Tween-20 0.5%, Triton-100 1.2% + PI solution). Samples were vortexed and incubated 15 min on ice. 600µL of Dilution buffer (100mM NaCl, 1mM EDTA, 25mM TRIS pH 7.5, 1mM DTT, Tween-20 0.5%, + PI solution) was then added and samples were centrifuged at 16.000 g for 30 min at 4°C. The resulting supernatant was pre-cleared by a 1 hour incubation at 4°C with 30µL Protein G beads equilibrated with wash buffer.

The cleared supernatants (inputs) were then incubated at 4°C with 30μL of the anti-Seh1, control, anti-Nup107- or anti-Nup85-coated Protein G beads. After overnight (for anti-Seh1) or 2 hours incubation (for anti-Nup107 and anti-Nup85), samples were centrifuged and washed 5 times in wash buffer. The proteins were either eluted in 40μL of Laemmli and boiled 10 minutes for subsequent western blot analysis (**Fig. 5B**), or split in 2 and then either eluted in 20μL of Laemmli and boiled 3 minutes for subsequent western blot analyses, or processed for both mass-spectrometry and western blot analyses (for experiments presented in **Fig. S3**).

#### Samples preparation prior to LC-MS/MS analysis

Proteins on beads were digested overnight at 37°C with trypsin (Promega, Madison, WI, USA) in a 25-mM NH_4_HCO_3_ buffer (0.2µg trypsin in 20µL). The resulting peptides were desalted using ZipTip μ-C18 Pipette Tips (Pierce Biotechnology, Rockford, IL, USA).

### LC-MS/MS acquisition

Samples were analyzed using an Orbitrap Fusion, coupled to a Nano-LC Proxeon 1200, equipped with an easy spray ion source (Thermo Scientific, Waltham, MA, USA). Peptides were loaded with an online preconcentration method and separated by chromatography using a Pepmap-RSLC C18 column (0.75 x 750 mm, 2 μm, 100 Å) from Thermo Scientific, equilibrated at 50°C and operated at a flow rate of 300 nl/min. Solvents (MS grade H_2_O, formic acid (FA) and Acetonitrile (ACN)) were from Thermo Chemical (Waltham, MA, USA).

Peptides were eluted by a gradient of solvent A (H_2_O, 0.1 % FA) and solvent B (ACN/H_2_O 80/20, 0.1% FA). The column was first equilibrated 5 min with 95 % of solvent A, then solvent B was raised to 28 % in 105 min and to 40% in 15 min. Finally, the column was washed with 95% solvent B during 20 min and re-equilibrated at 95% solvent A during 10 min. On the Orbitrap Fusion instrument, peptides precursor masses were analyzed in the Orbitrap cell in full ion scan mode, at a resolution of 120,000, a mass range of *m/z* 350-1550 and an AGC target of 4.10^5^. MS/MS were performed in the top speed 3s mode. Peptides were selected for fragmentation by Higher-energy C-trap Dissociation (HCD) with a Normalized Collisional Energy of 27% and a dynamic exclusion of 60 seconds. Fragment masses were measured in an Ion trap in the rapid mode, with and an AGC target of 1.10^4^. Monocharged peptides and unassigned charge states were excluded from the MS/MS acquisition. The maximum ion accumulation times were set to 100 ms for MS and 35 ms for MS/MS acquisitions respectively.

#### Data analysis

Raw data were processed on Proteome Discoverer 2.2 with the mascot node (Mascot version 2.5.1) and the Swissprot protein database release 2017_06. The *Mus musculus* taxonomy was used and a maximum of 2 missed cleavages was authorized. Precursor and fragment mass tolerances were set to 7 ppm and 0.5 Da. The following Post-translational modifications were included as variable: Acetyl (Protein N-term), Oxidation (M), Phosphorylation (STY). Spectra were filtered using a 1% FDR with the percolator node.

### Statistics

For cell confluence analyses, statistical analyses were performed at the latest time points (48h for cell growth in the pluripotent state and day 5 for neuroectodermal differentiation) using paired two-tailed Student’s t-test. For each mutant cell line, the % of confluence was compared to that of *WT* cells measured in the same experiment. To obtain more robust statistics, the paired two-tailed Student’s t-test was also used to compare all the values obtained with distinct clones bearing the same mutation to *WT* cells. For studies of interphase and mitosis duration and for quantifications of fluorescence intensity at the NE, statistical analyses were performed using unpaired non-parametric Mann-Whitney test. For nuclear surfaces, statistical analyses were performed using paired two-tailed Student’s t-test. P values and significance: ****: P <0.0001; ***: P <0.001; **: P <0.01; * :P <0.05.

## List of Symbols and Abbreviations used

mAID: mini Auxin Inducible Degron
mESC: mouse embryonic stem cell
NE: nuclear envelope
NPC: Nuclear pore complex
Nups: nucleoporins
n.s.: not significant
SD: standard deviation
*Tigre* locus: *Tightly regulated* locus
WT: Wild type.

## DATA AND SOFTWARE AVAILABILITY

The mass spectrometry proteomics data reported in this study have been deposited to the ProteomeXchange Consortium (http://proteomecentral.proteomexchange.org) via the PRIDE partner repository (Perez-Riverol et al., 2019) with the dataset identifier PXD022190.

The original 16-bit images and montages of the western blots used in this study are available as Mendeley dataset under http://dx.doi.org/10.17632/8g59mp92bs.1

## Acknowledgments

We are grateful to V. Heyer for generating the CRISPR/Cas9 vectors, to B. Souquet and A. Berto for advices regarding mESCs culture, and to P. Navarro and N. Festuccia for advices and reagents regarding integration at the *Tigre* locus and degron approaches. We kindly thank Jan Kosinski for discussions about the Nup85-Seh1 model, D. Forbes for sharing the Nup85 antibodies, N. Minc and S. Dmitrieff for their advices in the analysis of nuclear size, Liang Zhang for advise regarding anti-Seh1 immunoprecipitation experiments, and C. Boumendil and R. Karess for critical reading of the manuscript. We thank the proteomics core facility at the Institut Jacques-Monod, notably C. Garcia and L. Lignières, for the LC-MS/MS experiments. We also acknowledge the ImagoSeine core facility of the Institut Jacques Monod, notably M. Fradet and X. Baudin for help with cell sorting and spinning disk imaging, respectively.

## Competing interests

No competing interests declared.

## Funding

Work in the laboratory of VD was supported by the Centre national de la recherche scientifique (CNRS), the “Fondation pour la Recherche Médicale” (Foundation for Medical Research) under grant No DEQ20150734355, “Equipe FRM 2015” to VD and by the Labex Who Am I? (ANR-11-LABX-0071; Idex ANR-11-IDEX-0005-02). AG received a PhD fellowship from the INSPIRE, H2020-MSCA-COFUND-2014, Marie Skłodowska-Curie Co-funding of regional, national and international programmes, grant agreement No 665850, and a “transition post-doc grant from the Labex Who Am I?, AV received a post-doc grant from the Labex Who Am I? and CO a fellowship from Ecole Doctorale BioSPC, Université de Paris. Part of the LC-MS/MS equipment was founded by the Region Ile-de-France (SESAME 2013 Q-Prot-B&M - LS093471), the Paris-Diderot University (ARS 2014-2018), and CNRS (Moyens d’Equipement Exceptionnel INSB 2015). The ImagoSeine core facility was supported by founds from IBISA and the France-Bioimaging (ANR-10-INBS-04) infrastructures.

## AUTHOR CONTRIBUTIONS

A.G-E., A.V., C.O., B.R-S-M., and V.D. conceived and designed the experiments. A.G-E., A.V., and C.O. performed the experiments. A.G-E., A.V., C.O., and V.D. analyzed the data. A.G-E., A.V., and V.D. wrote the manuscript with contribution from all co-authors.

## Inventory of Supplementary Material

### Supplemental Figures and Figure legend

**Figure S1, related to Fig. 1.**
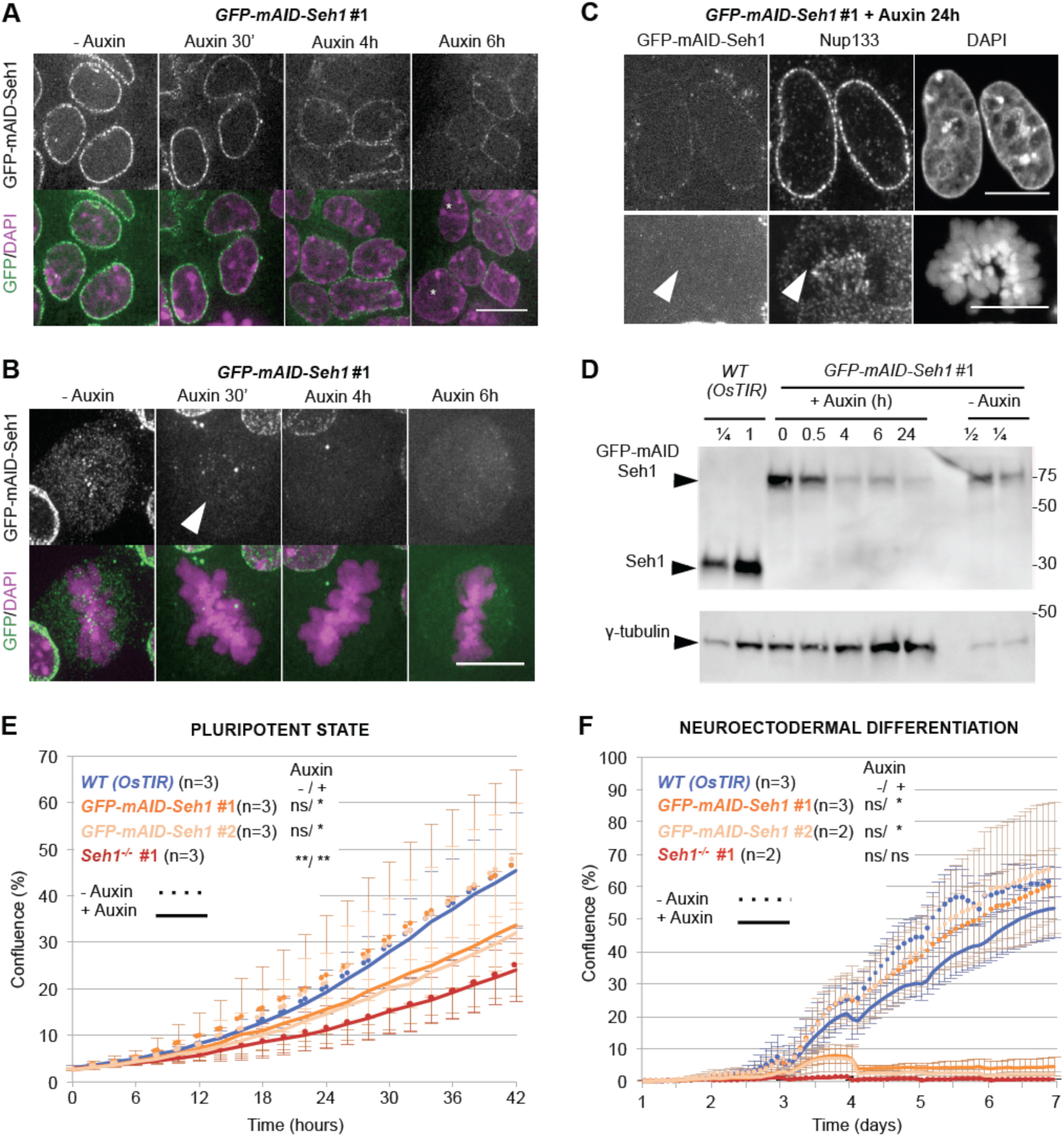
Auxin-induced GFP-mAID-Seh1 degradation in mESCs recapitulates *Seh1^-/-^* phenotypes. **A-B.** GFP-mAID-Seh1 localizes at NE in interphase cells (**A**; a single confocal plane is presented) and is enriched at kinetochores in mitotic cells (**B**; a projection of 9 optical sections is presented). After 30 minutes of auxin treatment, the GFP signal is only slightly decreased at the NE in interphase cells but no longer detectable in mitotic cells (arrowhead). Longer treatments (4 to 6 hours) with auxin lead to a progressive decrease of the NE signal, and to the appearance of GFP negative cells (*, likely corresponding to cells that went through mitosis). Scale bars, 10 µm. **C.** Localization of Nup133 in interphase (upper panels) and mitotic (lower panels) *GFP-mAID-Seh1* #1 mESCs after 24h of auxin treatment. The arrowheads point to the kinetochores that are labelled by Nup133 despite the lack of GFP-mAID-Seh1. Scale bar, 10 µm. **D.** Whole cell extracts from *WT-OsTIR* and *GFP-mAID-Seh1 #1* mESCs, treated or not with auxin for the indicated time, were analyzed by western blot using anti-Seh1 (top) and an anti-gamma-tubulin antibodies (bottom, used as loading control). 1/2 and 1/4 dilutions of *WT (OsTIR)* and non-treated *GFP-mAID-Seh1* #1 cell extracts were also loaded. Molecular weights are indicated (kilodaltons). **E.** Cell growth at pluripotent stage of the indicated cell lines was analyzed using cell confluence measurements with the IncuCyte→ system. Cells were treated with Auxin for 18 to 26 h prior to time 0. Error bars correspond to standard deviations from 3 independent experiments. **F.** The % of cell confluence upon neuroectodermal differentiation was quantified with the IncuCyte→ system. Cells were exposed to Auxin at the beginning of differentiation (day 0). Error bars correspond to standard deviations of 4 (*GFP-mAID-Seh1* #1 ad *OsTIR* WT) or 3 (*GFP-mAID-Seh1* #2 and *Seh1^-/^* #1) distinct wells acquired in 3 and 2 independent experiments, respectively. Statistical analyses were performed using the two-tailed Paired Student’s t-test (See Materials and Methods), by comparing all control (-Auxin) or treated (+Auxin) mutant cell lines to *Wt (OsTIR)* -Auxin or +Auxin, respectively.

**Figure S2, related to Fig. 2:**
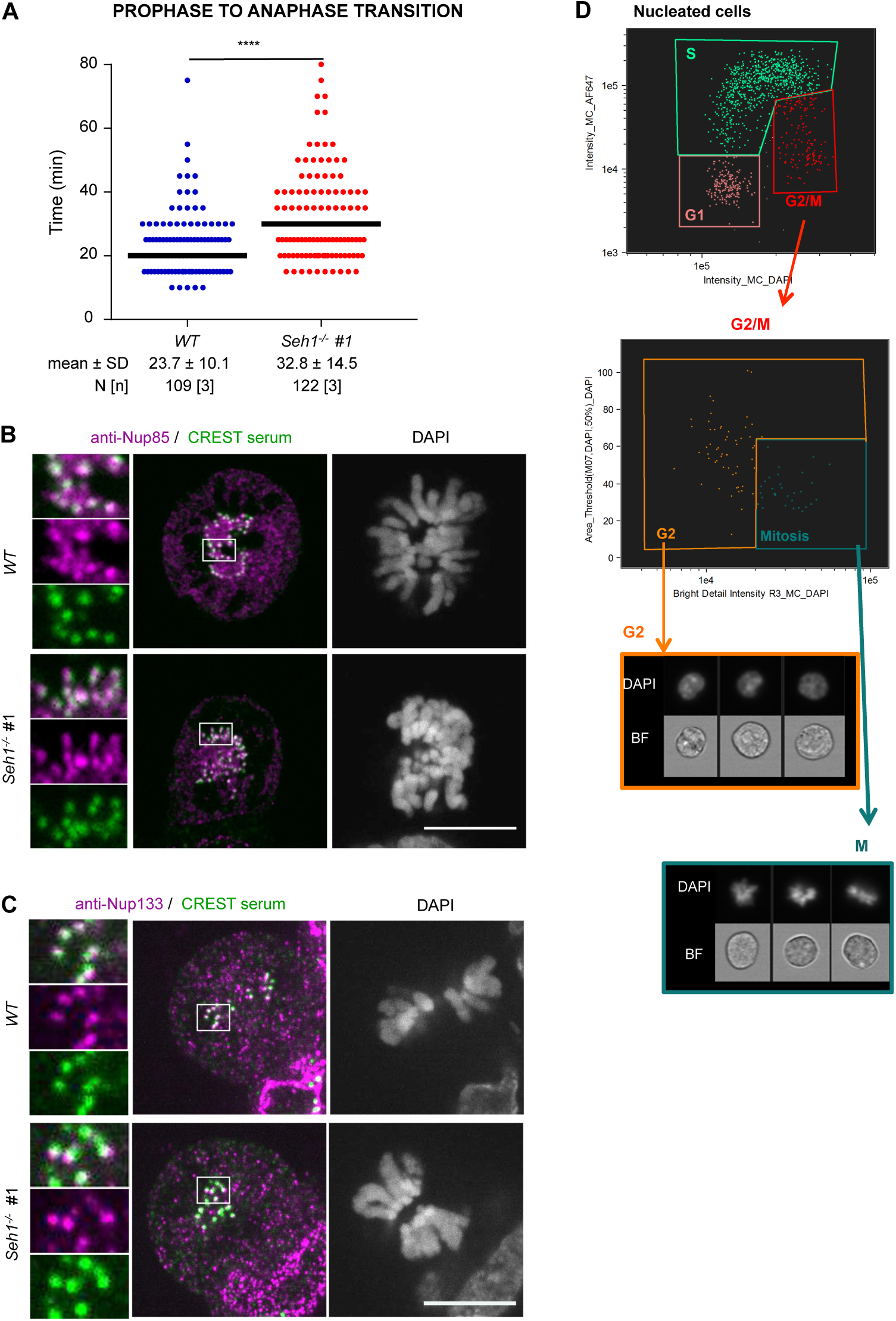
*Seh1^-/-^* mESCs have prolonged mitosis despite proper localization of the Y-complex at kinetochores. **A.** Quantification of the time spent from prometaphase to anaphase in *WT* and *Seh1^-/-^* mESCs. Each dot represents one cell and the median time is represented with a black bar. The number (N) of cells, quantified in 3 independent experiments is indicated**. B, C.** Representative spinning-disk images of mitotic *WT* and *Seh1^-/-^* mESCs, immunolabeled with (**C**) anti-Nup85 or (**D**) anti-Nup133, along with CREST serum (centromere marker) and stained with DAPI. A projection of three Z sections is presented. 3-fold enlargements of the boxed areas are also presented. Scale bars, 10 µm. **D.** Representative plots of the last steps of our ImageStream→ cell cycle analyses (*WT* mESCs). Top: the G1, S and G2/M phases of the cell cycle were defined based on DNA content (DAPI intensity) and EdU incorporation (AF647 channel). Middle: the G2 and M (Mitotic) population were then discriminated based on DAPI intensity and area. Bottom: bright Field (BF) and corresponding DAPI images of representative cells defined by these gating as G2 (orange) or mitotic (blue) are presented.

**Figure S3, related to Figs 6-8.**
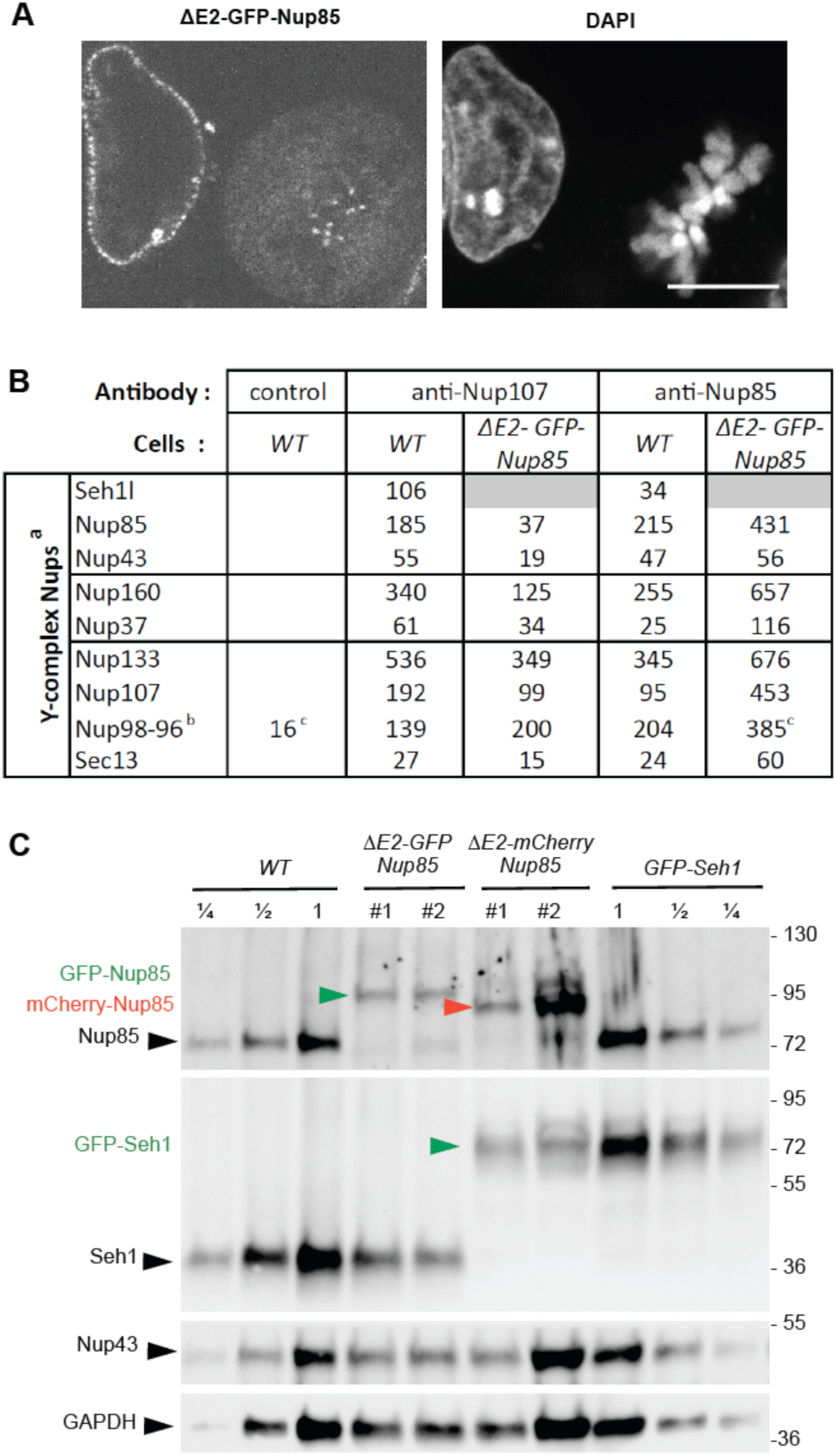
Characterization of the *ΔE2-GFP-Nup85* and *ΔE2-mCherry-Nup85* cell lines. (**A**) Representative spinning disk images (single plane) of interphase (left) and mitotic (right) *ΔE2-GFP-Nup85* mESCs fixed and stained with DAPI. Scale bar, 10 μm. **B**. The table provides the Mascot score for an immunoprecipitation experiment (see Materials and Methods) using either a pre-immune serum (control) or anti-Nup107 or -Nup85 antibodies, to pull down the Y-complex from *WT* or *ΔE2-GFP-Nup85* mESC protein extracts. ^a^Elys/AHCTF1 was not identified in this immunoprecipitation experiment. ^b^The autoproteolytic clivage of the Nup98-Nup96 precursor generates Nup98 and Nup96 (Fontoura et al., 1999); only the latter is a stable component of the Y-complex. Nearly all (35 out of 36) peptides identified in these immune pellets originates from Nup96. ^C^ Samples in which the Nup98 peptide was identified. **C.** Whole cell extracts of the indicated cell lines were analyzed by western-blot using anti-Nup85, -Seh1, -Nup43, and -GAPDH antibodies. Two-and four fold dilution (1/2, 1/4) of the *WT* and *GFP-Seh1* mESC extracts were also loaded. Molecular markers are indicated on the right (kDa).

**Figure S4, related to Materials and Methods and Tables S2 and S3:**
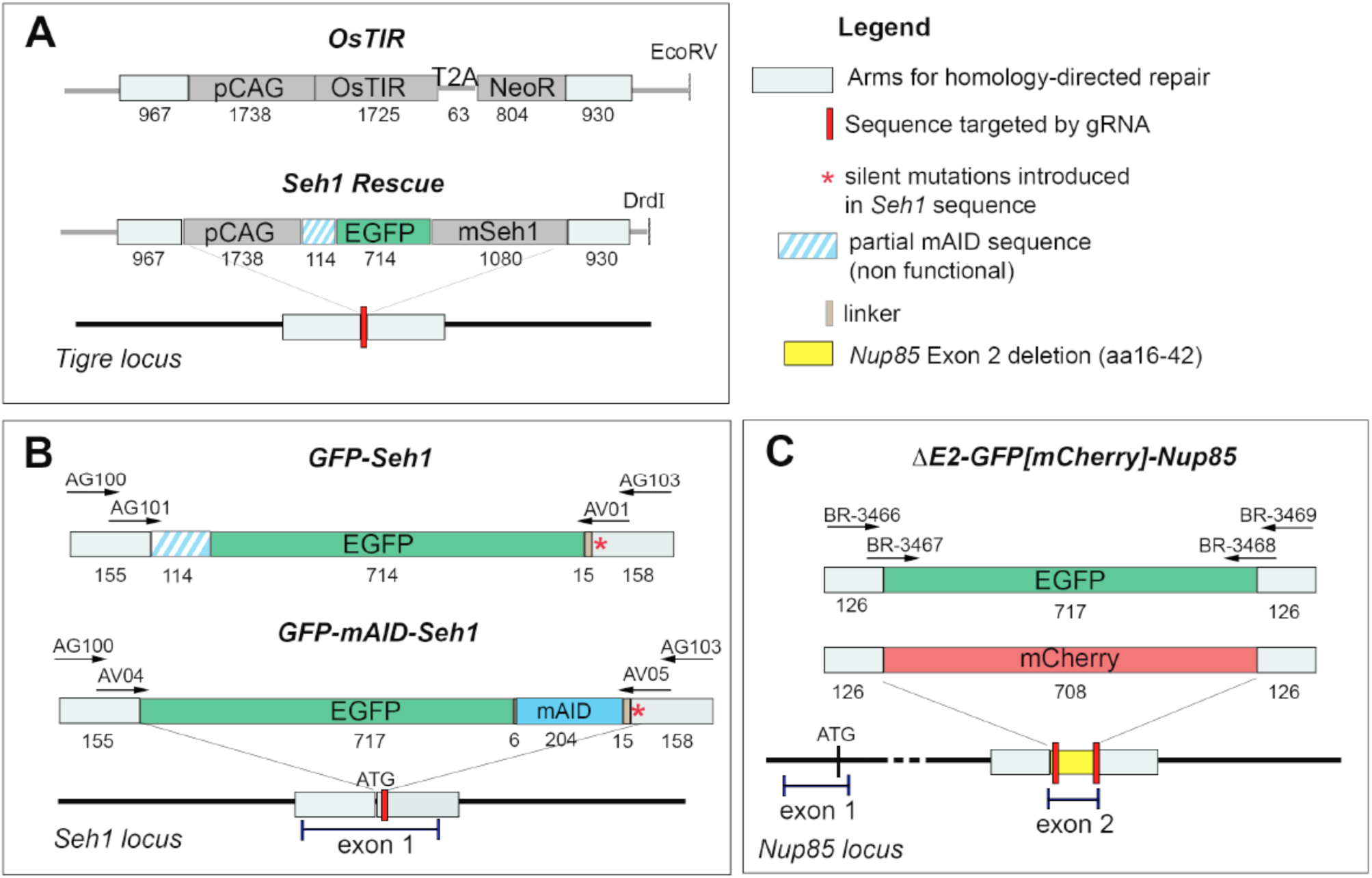
Strategies for CRISPR-Cas9-mediated cell line establishment via homologous recombination. The constructs used for homologous recombination (**A**, plasmid digested with the indicated restriction enzyme, and **B-C**, PCR products: the sequences of the oligonucleotides (arrows) used to amplify the template are listed in **Table S3**) leading to the indicated cell lines are represented above each targeted genomic locus (**A**: *Tigre*, B: *Seh1*, and **C**: *Nup85*). The sequences targeted by the gRNAs (red boxes), the sequences used for homology-directed repair (light grey boxes), and when relevant, the position of the exons and the first ATG are displayed. The size of the various segments is indicated in bp. Note that the construct used for GFP-Seh1 expression (A. *Seh1 rescue* and B. *GFP-Seh1* lines) further contains a non-functional 38 amino-acid long fragment of mAID. In **B.**, the red star indicates the position of silent CRISPR/Cas9 blocking mutations introduced in one Seh1 HR-arm.

### Supplemental Tables

**Table S1:** Plasmids.

| Plasmids used (sequence upon request) | Source | Identifier |
| --- | --- | --- |
| pX-280_Cas9WT-EGFP-Seh1gRNA1 - Seh1gRNA 2 | This paper | #1808 |
| pU6-sgTigre_CBh-Cas9-T2A-mCherry-3UTR | From P. Navarro Gil and N. Festuccia. (Festuccia et al., 2019) | #2061 |
| *Tigre* HR-pCAG-EGFP-mSeh1-*Tigre* HR | This paper | #2077 |
| *Tigre* HR-pCAG-OsTir-T2A-NeoR-*Tigre* HR | From P. Navarro Gil and N. Festuccia. (Festuccia et al., 2019) | #2064 |
| pX-853_Cas9WTmCherry Seh1-gRNA1 | This paper | #2098 |
| pCMV-EGFP-mAID-hSeh1 | This paper | #2100 |
| GeneArt CRISPR Nuclease Vector (OFP_Cas9) | Invitrogen, Carlsbad, CA | A21174 |
| GeneArt CRISPR pOFP-Cas9-Mios gRNA2 | This paper | #2110 |
| GeneArt CRISPR pOFP-Cas9-Mios gRNA4 | This paper | #2111 |
| GeneArt CRISPR pOFP-Cas9-Nup43 gRNA1 | This paper | #2104 |
| pX-672-Cas9HF-mCherry-gRNANup85-1_ gRNANup85-2 | This paper | #2005 |
| pX-864-Cas9HF-GFP-gRNANup85-1_ gRNANup85-2 | This paper | #2101 |
| pBOS-H2B-Cherry-IRES-Neo | From Doye lab (Bolhy et al., 2011) | #824 |
| pBOS-H2B-GFP-IRES-Neo | From Doye lab (Bolhy et al., 2011) | #805 |

**Table S2, related to Experimental Procedures.**
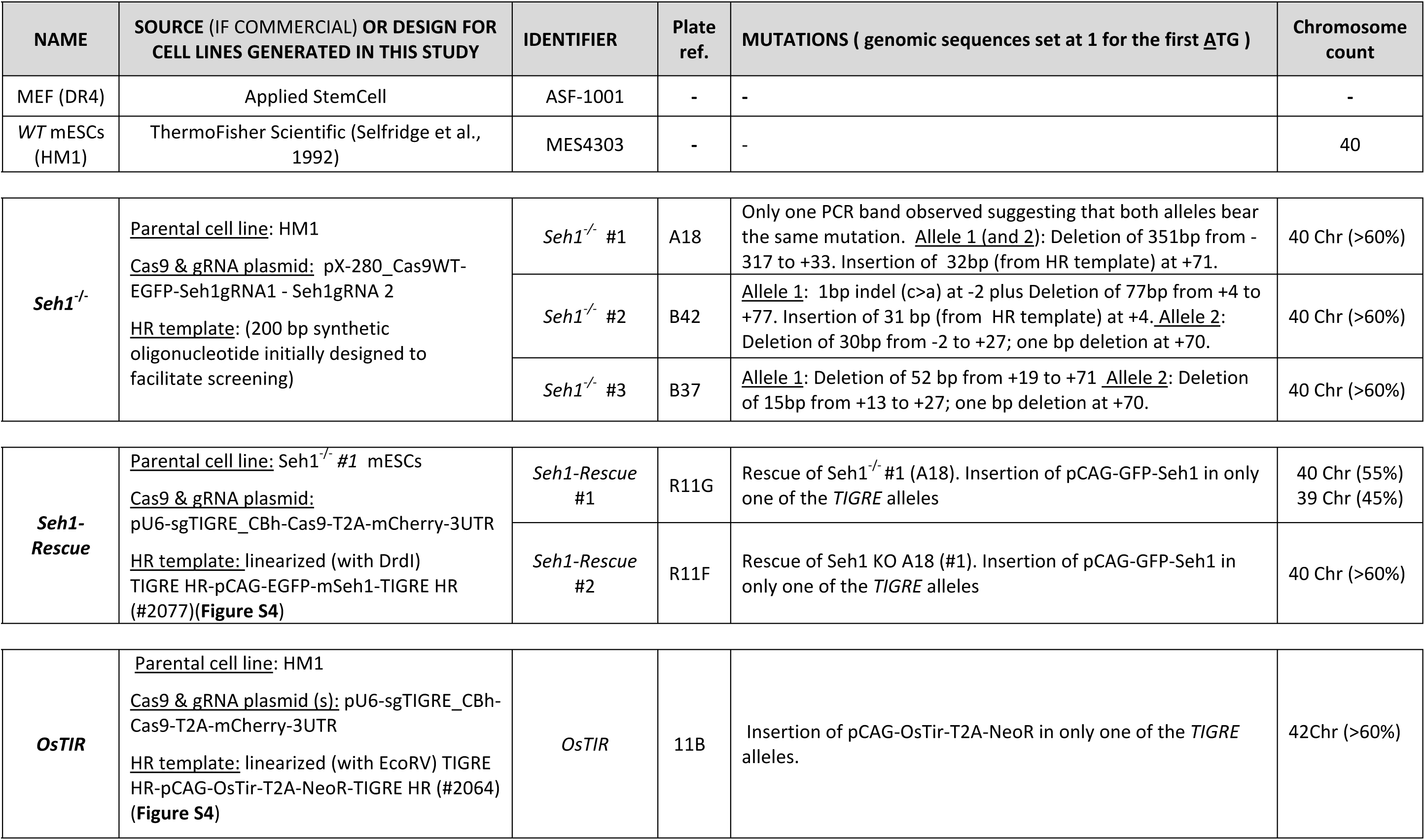
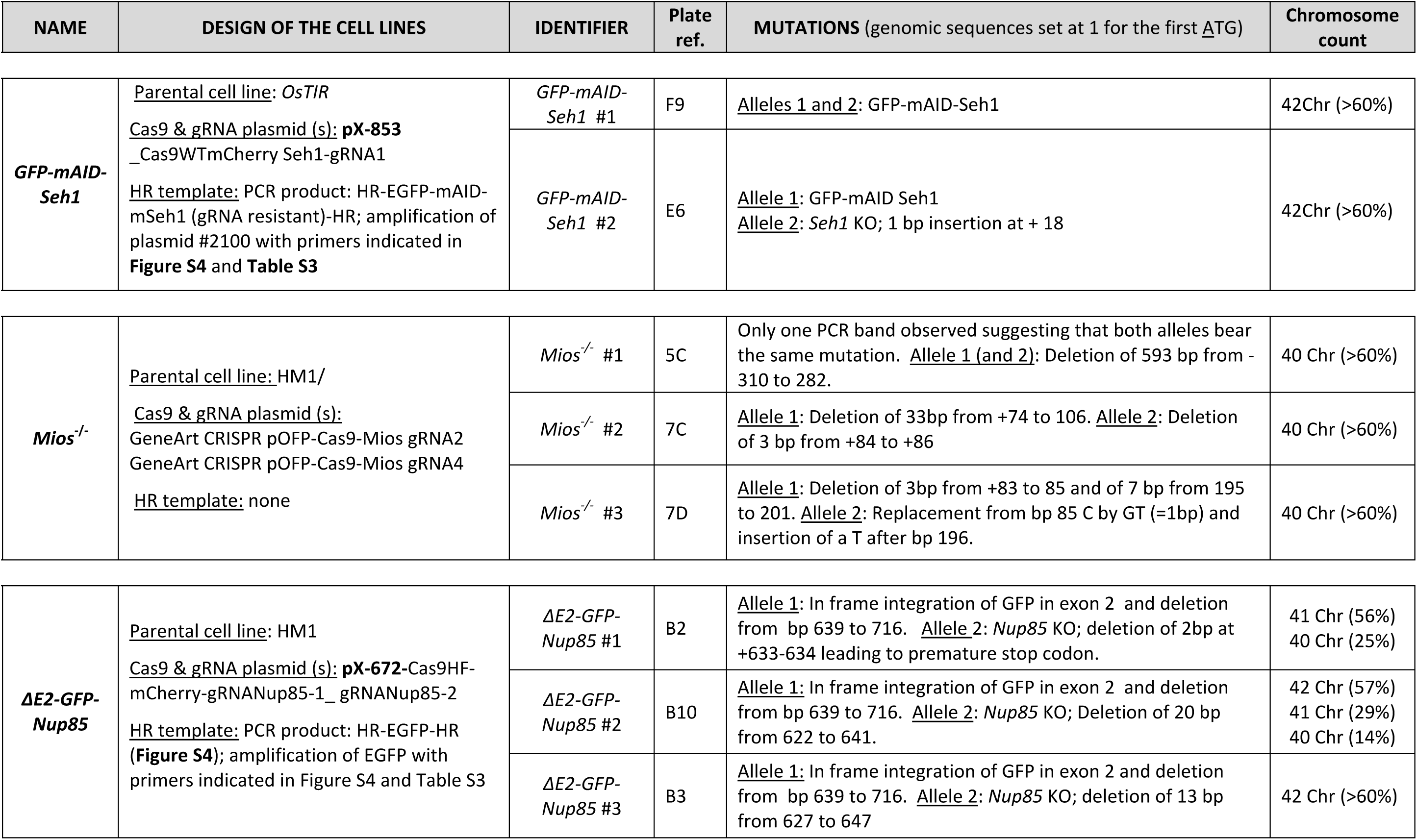
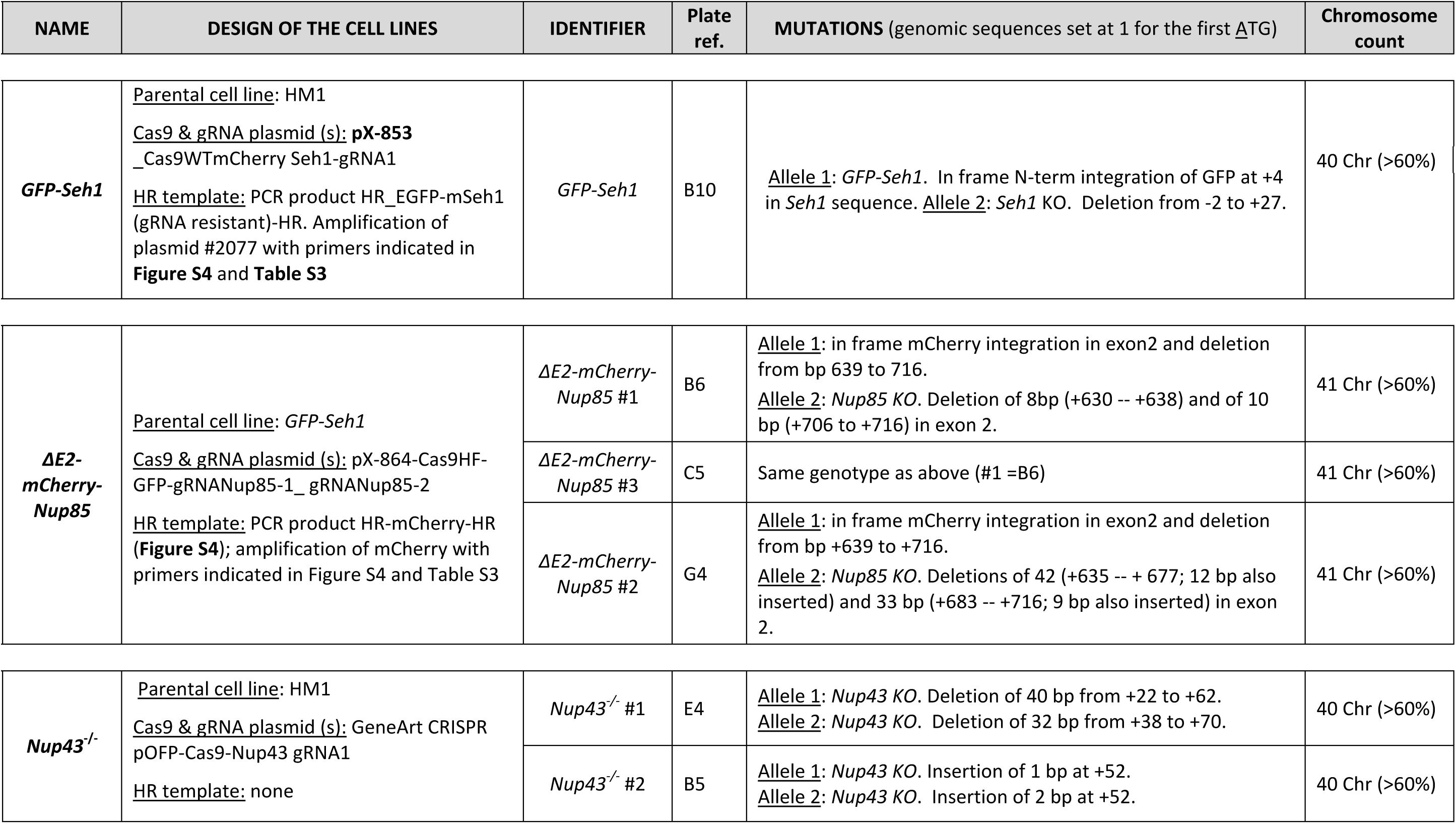
Cell lines used in this study

| NAME | SOURCE (IF COMMERCIAL) OR DESIGN FOR CELL LINES GENERATED IN THIS STUDY | IDENTIFIER | Plate ref. | MUTATIONS ( genomic sequences set at 1 for the first <u>A</u>TG ) | Chromosome count |
| --- | --- | --- | --- | --- | --- |
| MEF (DR4) | Applied StemCell | ASF-1001 | - | - | - |
| WT mESCs (HM1) | ThermoFisher Scientific (Selfridge et al., 1992) | MES4303 | - | - | 40 |
| **Seh1<sup>-/-</sup>** | <u>Parental cell line:</u> HM1<br><u>Cas9 & gRNA plasmid:</u> pX-280_Cas9WT-EGFP-Seh1gRNA1 - Seh1gRNA 2<br><u>HR template:</u> (200 bp synthetic oligonucleotide initially designed to facilitate screening) | *Seh1<sup>-/-</sup>* #1 | A18 | Only one PCR band observed suggesting that both alleles bear the same mutation. <u>Allele 1</u> (and <u>2</u>): Deletion of 351bp from -317 to +33. Insertion of 32bp (from HR template) at +71. | 40 Chr (>60%) |
|  |  | *Seh1<sup>-/-</sup>* #2 | B42 | <u>Allele 1</u>: 1bp indel (c>a) at -2 plus Deletion of 77bp from +4 to +77. Insertion of 31 bp (from HR template) at +4. <u>Allele 2</u>: Deletion of 30bp from -2 to +27; one bp deletion at +70. | 40 Chr (>60%) |
|  |  | *Seh1<sup>-/-</sup>* #3 | B37 | <u>Allele 1</u>: Deletion of 52 bp from +19 to +71 <u>Allele 2</u>: Deletion of 15bp from +13 to +27; one bp deletion at +70. | 40 Chr (>60%) |
| **Seh1-Rescue** | <u>Parental cell line:</u> Seh1<sup>-/-</sup> #1 mESCs<br><u>Cas9 & gRNA plasmid:</u> pU6-sgTIGRE_CBh-Cas9-T2A-mCherry-3UTR<br><u>HR template:</u> linearized (with DrdI) TIGRE HR-pCAG-EGFP-mSeh1-TIGRE HR (#2077)(**Figure S4**) | *Seh1-Rescue* #1 | R11G | Rescue of Seh1<sup>-/-</sup> #1 (A18). Insertion of pCAG-GFP-Seh1 in only one of the *TIGRE* alleles | 40 Chr (55%)<br>39 Chr (45%) |
|  |  | *Seh1-Rescue* #2 | R11F | Rescue of Seh1 KO A18 (#1). Insertion of pCAG-GFP-Seh1 in only one of the *TIGRE* alleles | 40 Chr (>60%) |
| **OsTIR** | <u>Parental cell line:</u> HM1<br><u>Cas9 & gRNA plasmid (s):</u> pU6-sgTIGRE_CBh-Cas9-T2A-mCherry-3UTR<br><u>HR template:</u> linearized (with EcoRV) TIGRE HR-pCAG-OsTir-T2A-NeoR-TIGRE HR (#2064) (**Figure S4**) | *OsTIR* | 11B | Insertion of pCAG-OsTir-T2A-NeoR in only one of the *TIGRE* alleles. | 42Chr (>60%) |

**Table S2- continued**
| NAME | DESIGN OF THE CELL LINES | IDENTIFIER | Plate ref. | MUTATIONS (genomic sequences set at 1 for the first ATG) | Chromosome count |
| --- | --- | --- | --- | --- | --- |
| **GFP-mAID-**<br>**Seh1** | <u>Parental cell line:</u> *OsTIR*<br><u>Cas9 & gRNA plasmid (s):</u> **pX-853**<br>_Cas9WTmCherry Seh1-gRNA1<br><u>HR template:</u> PCR product: HR-EGFP-mAID-mSeh1 (gRNA resistant)-HR; amplification of plasmid #2100 with primers indicated in **Figure S4** and **Table S3** | *GFP-mAID-*<br>*Seh1* #1 | F9 | <u>Alleles 1 and 2:</u> GFP-mAID-Seh1 | 42Chr (>60%) |
|  |  | *GFP-mAID-*<br>*Seh1* #2 | E6 | <u>Allele 1:</u> GFP-mAID Seh1<br><u>Allele 2:</u> *Seh1* KO; 1 bp insertion at + 18 | 42Chr (>60%) |
| **Mios<sup>-/-</sup>** | <u>Parental cell line:</u> HM1/<br><u>Cas9 & gRNA plasmid (s):</u><br>GeneArt CRISPR pOFP-Cas9-Mios gRNA2<br>GeneArt CRISPR pOFP-Cas9-Mios gRNA4<br><u>HR template:</u> none | *Mios<sup>-/-</sup>* #1 | 5C | Only one PCR band observed suggesting that both alleles bear the same mutation. <u>Allele 1 (and 2):</u> Deletion of 593 bp from - 310 to 282. | 40 Chr (>60%) |
|  |  | *Mios<sup>-/-</sup>* #2 | 7C | <u>Allele 1:</u> Deletion of 33bp from +74 to 106. <u>Allele 2:</u> Deletion of 3 bp from +84 to +86 | 40 Chr (>60%) |
|  |  | *Mios<sup>-/-</sup>* #3 | 7D | <u>Allele 1:</u> Deletion of 3bp from +83 to 85 and of 7 bp from 195 to 201. <u>Allele 2:</u> Replacement from bp 85 C by GT (=1bp) and insertion of a T after bp 196. | 40 Chr (>60%) |
| **ΔE2-GFP-**<br>**Nup85** | <u>Parental cell line:</u> HM1<br><u>Cas9 & gRNA plasmid (s):</u> **pX-672**-Cas9HF-mCherry-gRNANup85-1_ gRNANup85-2<br><u>HR template:</u> PCR product: HR-EGFP-HR (**Figure S4**); amplification of EGFP with primers indicated in Figure S4 and Table S3 | *ΔE2-GFP-*<br>*Nup85* #1 | B2 | <u>Allele 1:</u> In frame integration of GFP in exon 2 and deletion from bp 639 to 716. <u>Allele 2:</u> *Nup85* KO; deletion of 2bp at +633-634 leading to premature stop codon. | 41 Chr (56%)<br>40 Chr (25%) |
|  |  | *ΔE2-GFP-*<br>*Nup85* #2 | B10 | <u>Allele 1:</u> In frame integration of GFP in exon 2 and deletion from bp 639 to 716. <u>Allele 2:</u> *Nup85* KO; Deletion of 20 bp from 622 to 641. | 42 Chr (57%)<br>41 Chr (29%)<br>40 Chr (14%) |
|  |  | *ΔE2-GFP-*<br>*Nup85* #3 | B3 | <u>Allele 1:</u> In frame integration of GFP in exon 2 and deletion from bp 639 to 716. <u>Allele 2:</u> *Nup85* KO; deletion of 13 bp from 627 to 647 | 42 Chr (>60%) |

Table S2- continued
| NAME | DESIGN OF THE CELL LINES | IDENTIFIER | Plate ref. | MUTATIONS (genomic sequences set at 1 for the first ATG) | Chromosome count |
| --- | --- | --- | --- | --- | --- |
| **GFP-Seh1** | Parental cell line: HM1<br>Cas9 & gRNA plasmid (s): **pX-853**<br>_Cas9WTmCherry Seh1-gRNA1<br>HR template: PCR product HR_EGFP-mSeh1 (gRNA resistant)-HR. Amplification of plasmid #2077 with primers indicated in **Figure S4** and **Table S3** | *GFP-Seh1* | B10 | <u>Allele 1</u>: *GFP-Seh1*. In frame N-term integration of GFP at +4 in *Seh1* sequence. <u>Allele 2</u>: *Seh1* KO. Deletion from -2 to +27. | 40 Chr (>60%) |
| **$\Delta E2$-mCherry-Nup85** | Parental cell line: *GFP-Seh1*<br>Cas9 & gRNA plasmid (s): pX-864-Cas9HF-GFP-gRNANup85-1_gRNANup85-2<br>HR template: PCR product HR-mCherry-HR (**Figure S4**); amplification of mCherry with primers indicated in Figure S4 and Table S3 | *$\Delta E2$-mCherry-Nup85 #1* | B6 | <u>Allele 1</u>: in frame mCherry integration in exon2 and deletion from bp 639 to 716.<br><u>Allele 2</u>: *Nup85* KO. Deletion of 8bp (+630 -- +638) and of 10 bp (+706 to +716) in exon 2. | 41 Chr (>60%) |
|  |  | *$\Delta E2$-mCherry-Nup85 #3* | C5 | Same genotype as above (#1 =B6) | 41 Chr (>60%) |
|  |  | *$\Delta E2$-mCherry-Nup85 #2* | G4 | <u>Allele 1</u>: in frame mCherry integration in exon2 and deletion from bp +639 to +716.<br><u>Allele 2</u>: *Nup85* KO. Deletions of 42 (+635 -- + 677; 12 bp also inserted) and 33 bp (+683 -- +716; 9 bp also inserted) in exon 2. | 41 Chr (>60%) |
| ***Nup43*<sup>-/-</sup>** | Parental cell line: HM1<br>Cas9 & gRNA plasmid (s): GeneArt CRISPR pOFP-Cas9-Nup43 gRNA1<br>HR template: none | *Nup43*<sup>-/-</sup> #1 | E4 | <u>Allele 1</u>: *Nup43* KO. Deletion of 40 bp from +22 to +62.<br><u>Allele 2</u>: *Nup43* KO. Deletion of 32 bp from +38 to +70. | 40 Chr (>60%) |
|  |  | *Nup43*<sup>-/-</sup> #2 | B5 | <u>Allele 1</u>: *Nup43* KO. Insertion of 1 bp at +52.<br><u>Allele 2</u>: *Nup43* KO. Insertion of 2 bp at +52. | 40 Chr (>60%) |

**Table S3:**
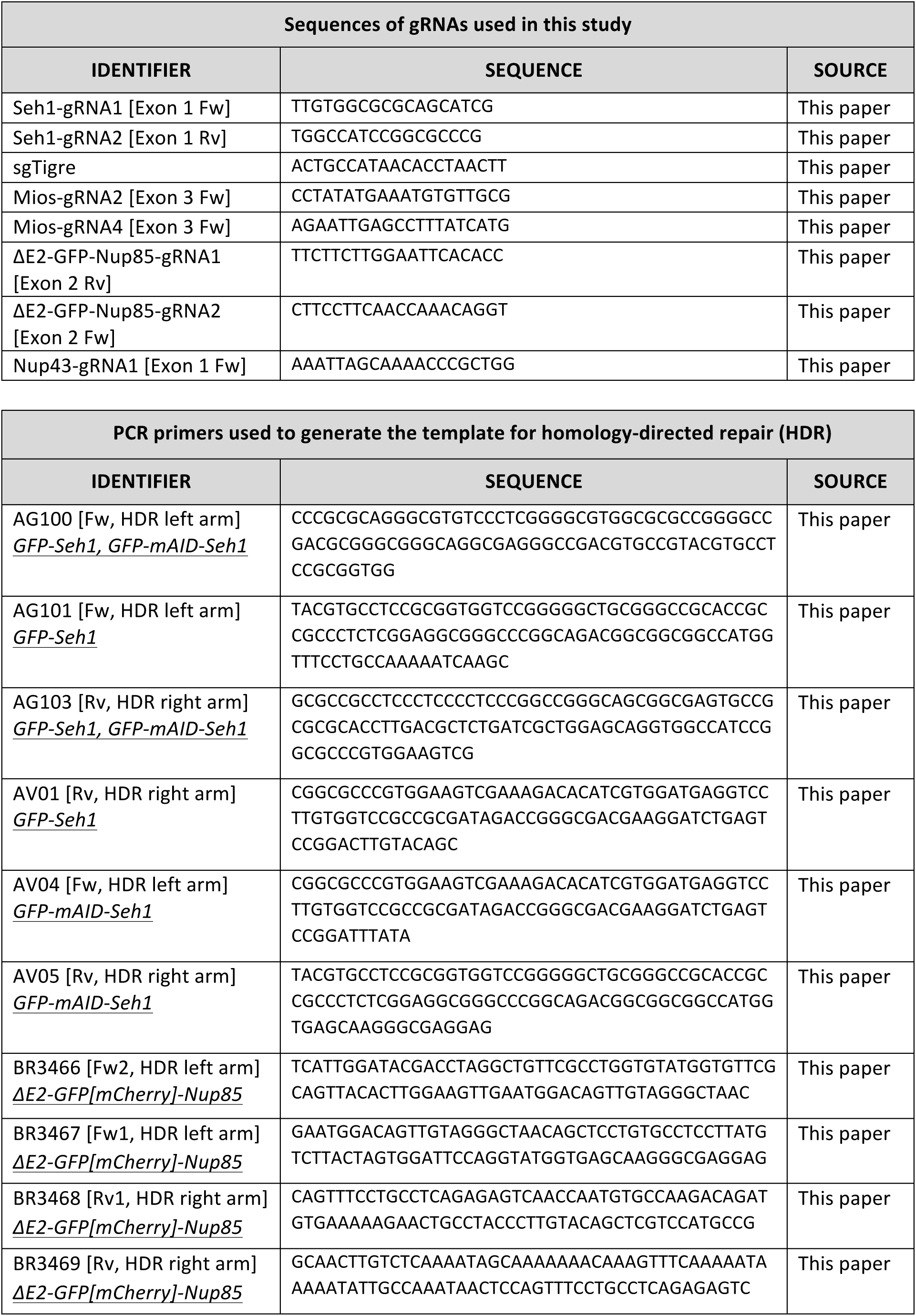
Sequences of gRNAs and PCR primers used to generate the template for homology-directed repair (HDR)

**Table S4:** Antibodies

| Primary antibodies | Dilution | Source | Identifier |
| --- | --- | --- | --- |
| Human autoantibody against-centromeres (CREST serum) | IF: 1/500 | ImmunoVision, Springdale, AR | HCT-0100 |
| Rabbit polyclonal anti-GAPDH | WB: 1/2000 | Trevigen, Minneapolis, MN | 2275-PC100 |
| Mouse anti-gamma-tubulin | WB: 1/50000 | Abcam, San Francisco, CA | Ab11316 |
| Rabbit polyclonal antibody anti-Mios | WB: 1/1000 | Cell signalling, Danvers, MA | #13557 |
| Affinity-purified rabbit polyclonal anti-Nup43 antibody | WB: 1/1000 | From Doye lab (Zuccolo et al., 2007) | # 571/08-111-C1 |
| Rabbit polyclonal anti-mNup85 serum | WB: 1/2000<br>IF: 1:1000<br>IP: 1/10 | From D. Forbes (Harel et al., 2003) | N/A |
| Rabbit monoclonal antibody anti-Nup98 C39A3 | IF: 1/20 | Cell signalling | #2598 |
| Rabbit polyclonal anti-Nup107 | WB: 1/1000<br>IP: 1/10 | From Doye lab (Belgareh et al., 2001) | #520-82 |
| Rat monoclonal anti-mouse Nup133 antibody #74; clone 9C2H8) | IF: 1/100 | From Doye lab (Berto et al., 2018) | N/A |
| Rabbit polyclonal antibody anti-Seh1 | WB: 1/1000<br>IP: 1/500 | Abcam, San Francisco, CA | ab218531 |
| Rabbit polyclonal antibody anti-Tpr | IF: 1/200 | Abcam, San Francisco, CA | ab84516 |
| Rabbit polyclonal antibody anti Ostir | WB: 1/1000 | From M.T. Kanemaki (Natsume et al., 2016) | N/A |
| Rabbit polyclonal antibody anti- Wdr24 | WB: 1/1000 | Proteintech, Manchester, UK | 20778-1-AP |

| Secondary antibodies | Dilution | Source | Identifier |
| --- | --- | --- | --- |
| Cy™3 AffiniPure Donkey anti-rabbit | IF: 1/1000 | Jackson ImmunoRes, West Grove, PA | 711-165-152 |
| Cy™3 AffiniPure Donkey anti-rat | IF: 1/1000 | Jackson ImmunoRes | 712-165-153 |
| Cy™3 AffiniPure Goat anti-rabbit | IF: 1/1000 | Jackson ImmunoRes | 111-165-144 |
| AlexaFluor 568 Donkey anti-rabbit | IF: 1/1000 | ThermoFisher Scientific | A10042 |
| Alexa 647 Donkey anti-human | IF: 1/500 | Jackson ImmunoRes | 709-605-149 |
| Cy™5 AffiniPure Donkey anti-rat | IF: 1/1000 | Jackson ImmunoRes | 712-175-153 |
| Peroxidase AffiniPure Donkey anti-rabbit | WB: 1/5000 | Jackson ImmunoRes. | 711-035-152 |
| Peroxidase AffiniPure Goat anti-mouse | WB: 1/5000 | Jackson ImmunoRes. | 115-035-068 |
| Peroxidase AffiniPure Goat anti-ratCy™5 AffiniPure Donkey Anti-Rat | WB: 1/5000 | Jackson ImmunoRes. | 112-035-167 |
| Rabbit TrueBlot ®: Anti-Rabbit IgG HRP | WB: 1/1000 | Rockland, Limerick, PA | 18-8816-33 |

**Supplemental Movies, related to Figure 1:**

Examples of neuroectodermal differentiation experiments as exported from IncuCyte→ device. Time and scale bars are indicated

- Movie S1: *WT*

- Movie S2: *Seh1 ^-/-^ #1*

- Movie S3: *Rescue #2*

## Notes

http://dx.doi.org/10.17632/8g59mp92bs.1

http://proteomecentral.proteomexchange.org/cgi/GetDataset?ID=PXD022190

### Competing Interest Statement

The authors have declared no competing interest.

### Summary of Updates

- Quantitative IF analyses on the Nup43-/- and ΔE2-GFP-Nup85 mutant cells with both Nup133 and Nup98 antibodies (in addition to TPR as analyzed in the previous version of the manuscript) confirm that all these Nups are similarly affected when the short arm of the Y-complex is altered. - Immunoprecipitation experiments now show that lack of Mios prevents Seh1 interaction with Wdr24, another GATOR2 complex component - We now provide statistical analysis of the confluency curves and quantitation of GFP-Seh1 kinetochore intensity in ΔE2-mCherry-Nup85 cells.

## References

Algret, R., Fernandez-Martinez, J., Shi, Y., Kim, S.J., Pellarin, R., Cimermancic, P., Cochet, E., Sali, A., Chait, B.T., Rout, M.P., et al. (2014). Molecular Architecture and Function of the SEA Complex, a Modulator of the TORC1 Pathway. Mol Cell Proteomics 13, 2855–2870.

Bar-Peled, L., Chantranupong, L., Cherniack, A.D., Chen, W.W., Ottina, K.A., Grabiner, B.C., Spear, E.D., Carter, S.L., Meyerson, M., and Sabatini, D.M. (2013). A Tumor suppressor complex with GAP activity for the Rag GTPases that signal amino acid sufficiency to mTORC1. Science 340, 1100–1106.

Brohawn, S.G., Leksa, N.C., Spear, E.D., Rajashankar, K.R., and Schwartz, T.U. (2008). Structural Evidence for Common Ancestry of the Nuclear Pore Complex and Vesicle Coats. Science 322, 1369–1373.

Buchwalter, A., Kaneshiro, J.M., and Hetzer, M.W. (2019). Coaching from the sidelines: the nuclear periphery in genome regulation. Nat Rev Genet 20, 39–50.

Cai, W., Wei, Y., Jarnik, M., Reich, J., and Lilly, M.A. (2016). The GATOR2 Component Wdr24 Regulates TORC1 Activity and Lysosome Function. PLoS Genet 12, e1006036.

Debler, E.W., Ma, Y., Seo, H.S., Hsia, K.C., Noriega, T.R., Blobel, G., and Hoelz, A. (2008). A Fence-like Coat for the Nuclear Pore Membrane. Mol Cell 32, 815–826.

Dokudovskaya, S., and Rout, M.P. (2015). SEA you later alli-GATOR - a dynamic regulator of the TORC1 stress response pathway. J Cell Sci 128, 2219–2228.

Doucet, C.M., Talamas, J.A., and Hetzer, M.W. (2010). Cell cycle-dependent differences in nuclear pore complex assembly in metazoa. Cell 141, 1030–1041.

Faria, A.M.C., Levay, A., Wang, Y., Kamphorst, A.O., Rosa, M.L.P., Nussenzveig, D.R., Balkan, W., Chook, Y.M., Levy, D.E., and Fontoura, B.M.A. (2006). The Nucleoporin Nup96 Is Required for Proper Expression of Interferon-Regulated Proteins and Functions. Immunity 24, 295–304.

Engler, C., Gruetzner, R., Kandzia, R., and Marillonnet, S. (2009). Golden Gate Shuffling: A One-Pot DNA Shuffling Method Based on Type IIs Restriction Enzymes. PLOS ONE 4, e5553.

Fernandez-Martinez, J., Kim, S.J., Shi, Y., Upla, P., Pellarin, R., Gagnon, M., Chemmama, I.E., Wang, J., Nudelman, I., Zhang, W., et al. (2016). Structure and Function of the Nuclear Pore Complex Cytoplasmic mRNA Export Platform. Cell 167, 1215–1228.e1225.

Hampoelz, B., Andres-Pons, A., Kastritis, P., and Beck, M. (2019). Structure and Assembly of the Nuclear Pore Complex. Annu Rev Biophys 48, 515–536.

Harel, A., Orjalo, A.V., Vincent, T., Lachish-Zalait, A., Vasu, S., Shah, S., Zimmerman, E., Elbaum, M., and Forbes, D.J. (2003). Removal of a single pore subcomplex results in vertebrate nuclei devoid of nuclear pores. Mol Cell 11, 853–864.

Hezwani, M., and Fahrenkrog, B. (2017). The functional versatility of the nuclear pore complex proteins. Semin Cell Dev Biol 68, 2–9.

Huang, G., Zhang, Y., Zhu, X., Zeng, C., Wang, Q., Zhou, Q., Tao, Q., Liu, M., Lei, J., Yan, C., and Shi, Y.(2020). Structure of the cytoplasmic ring of the Xenopus laevis nuclear pore complex by cryo-electron microscopy single particle analysis. Cell Res 30,520–531

Jevtić, P., Mukherjee, R.N., Chen, P., and Levy, D.L. (2019). Altering the levels of nuclear import factors in early Xenopus laevis embryos affects later development. PLoS ONE 14, e0215740.

Kosinski, J., Mosalaganti, S., von Appen, A., Teimer, R., DiGuilio, A.L., Wan, W., Bui, K.H., Hagen, W.J.H., Briggs, J.A.G., Glavy, J.S., et al. (2016). Molecular architecture of the inner ring scaffold of the human nuclear pore complex. Science 352, 363–365.

Lin, D.H., and Hoelz, A. (2019). The Structure of the Nuclear Pore Complex (An Update). Annu Rev Biochem 88, 725–783.

Liu, Z., Yan, M., Liang, Y., Liu, M., Zhang, K., Shao, D., Jiang, R., Li, L., Wang, C., Nussenzveig, D.R., et al. (2019). Nucleoporin Seh1 Interacts with Olig2/Brd7 to Promote Oligodendrocyte Differentiation and Myelination. Neuron 102, 587–601.e7

Loiodice, I., Alves, A., Rabut, G., Van Overbeek, M., Ellenberg, J., Sibarita, J.B., and Doye, V. (2004). The entire Nup107-160 complex, including three new members, is targeted as one entity to kinetochores in mitosis. Mol Biol Cell 15, 3333–3344.

Lupu, F., Alves, A., Anderson, K., Doye, V., and Lacy, E. (2008). Nuclear pore composition regulates neural stem/progenitor cell differentiation in the mouse embryo. Dev Cell 14, 831–842.

Machado, S., Mercier, V., and Chiaruttini, N. (2019). LimeSeg: a coarse-grained lipid membrane simulation for 3D image segmentation. BMC bioinformatics 20, 2.

Moreira, T.G., Zhang, L., Shaulov, L., Harel, A., Kuss, S.K., Williams, J., Shelton, J., Somatilaka, B., Seemann, J., Yang, J., et al. (2015). Sec13 Regulates Expression of Specific Immune Factors Involved in Inflammation In Vivo. Sci Rep 5, 17655.

Natsume, T., Kiyomitsu, T., Saga, Y., and Kanemaki, M.T. (2016). Rapid Protein Depletion in Human Cells by Auxin-Inducible Degron Tagging with Short Homology Donors. Cell Rep 15, 210–218.

Neumann, N., Lundin, D., and Poole, A.M. (2010). Comparative Genomic Evidence for a Complete Nuclear Pore Complex in the Last Eukaryotic Common Ancestor. PLoS ONE 5, e13241.

Okita, K., Kiyonari, H., Nobuhisa, I., Kimura, N., Aizawa, S., and Taga, T. (2004). Targeted disruption of the mouse ELYS gene results in embryonic death at peri-implantation development. Genes Cells 9, 1083–1091.

Pascual-Garcia, P., and Capelson, M. (2019). Nuclear pores in genome architecture and enhancer function. Curr Opin Cell Biol 58, 126–133.

Perez-Riverol, Y., Csordas, A., Bai, J., Bernal-Llinares, M., Hewapathirana, S., Kundu, D.J., Inuganti, A., Griss, J., Mayer, G., Eisenacher, M., et al. (2019). The PRIDE database and related tools and resources in 2019: improving support for quantification data. Nucleic Acids Res 47, D442–D450.

Platani, M., Samejima, I., Samejima, K., Kanemaki, M.T., and Earnshaw, W.C. (2018). Seh1 targets GATOR2 and Nup153 to mitotic chromosomes. J Cell Sci 131, jcs213140.

Platani, M., Santarella-Mellwig, R., Posch, M., Walczak, R., Swedlow, J.R., and Mattaj, I.W. (2009). The Nup107-160 nucleoporin complex promotes mitotic events via control of the localization state of the chromosome passenger complex. Mol Biol Cell 20, 5260–5275.

Platani, M., Trinkle-Mulcahy, L., Porter, M., Jeyaprakash, A.A., and Earnshaw, W.C. (2015). Mio depletion links mTOR regulation to Aurora A and Plk1 activation at mitotic centrosomes. J Cell Biol. 210, 45–62.

Rabut, G., Doye, V., and Ellenberg, J. (2004). Mapping the dynamic organization of the nuclear pore complex inside single living cells. Nat Cell Biol 6, 1114–1121.

Rasala, B.A., Orjalo, A.V., Shen, Z.X., Briggs, S., and Forbes, D.J. (2006). ELYS is a dual nucleoporin/kinetochore protein required for nuclear pore assembly and proper cell division. Proc Natl Acad Sci USA 103, 17801–17806.

Salama, N.R., Yeung, T., and Schekman, R.W. (1993). The Sec13p Complex and Reconstitution of Vesicle Budding from the Er with Purified Cytosolic Proteins. EMBO J 12, 4073–4082.

Scholz, B.A., Sumida, N., de Lima, C.D.M., Chachoua, I., Martino, M., Tzelepis, I., Nikoshkov, A., Zhao, H.L., Mehmood, R., Sifakis, E.G., et al. (2019). WNT signaling and AHCTF1 promote oncogenic MYC expression through super-enhancer-mediated gene gating. Nat Genet 51, 1723–1731.

Selfridge, J., Pow, A.M., McWhir, J., Magin, T.M., and Melton, D.W. (1992). Gene targeting using a mouse HPRT minigene/HPRT-deficient embryonic stem cell system: Inactivation of the mouseERCC-1 gene. Somat Cell Molec Gen 18, 325–336.

Senger, S., Csokmay, J., Tanveer, A., Jones, T.I., Sengupta, P., and Lilly, M.A. (2011). The nucleoporin Seh1 forms a complex with Mio and serves an essential tissue-specific function in Drosophila oogenesis. Development 138, 2133–2142.

Souquet, B., Freed, E., Berto, A., Andric, V., Auduge, N., Reina-San-Martin, B., Lacy, E., and Doye, V. (2018). Nup133 Is Required for Proper Nuclear Pore Basket Assembly and Dynamics in Embryonic Stem Cells. Cell Rep 23, 2443–2454.

Terashima, Y., Toda, E., Itakura, M., Otsuji, M., Yoshinaga, S., Okumura, K., Shand, F.H.W., Komohara, Y., Takeda, M., Kokubo, K., et al. (2020). Targeting FROUNT with disulfiram suppresses macrophage accumulation and its tumor-promoting properties. Nat Commun 11, 609.

Vollmer, B., Lorenz, M., Moreno-Andrés, D., Bodenhöfer, M., De Magistris, P., Astrinidis, S.A., Schooley, A., Flötenmeyer, M., Leptihn, S., and Antonin, W. (2015). Nup153 Recruits the Nup107-160 Complex to the Inner Nuclear Membrane for Interphasic Nuclear Pore Complex Assembly. Dev Cell 33, 717–728.

von Appen, A., Kosinski, J., Sparks, L., Ori, A., DiGuilio, A.L., Vollmer, B., Mackmull, M.T., Banterle, N., Parca, L., Kastritis, P., et al. (2015). In situ structural analysis of the human nuclear pore complex. Nature 526, 140–143.

Walther, T.C., Alves, A., Pickersgill, H., Loiodice, I., Hetzer, M., Galy, V., Hulsmann, B.B., Kocher, T., Wilm, M., Allen, T., et al. (2003). The conserved Nup107-160 complex is critical for nuclear pore complex assembly. Cell 113, 195–206.

Webster, B.M., and Lusk, C.P. (2016). Border safety: quality control at the nuclear envelope. Trends Cell Biol 26, 29–39.

Ying, Q.L., and Smith, A.G. (2003). Defined conditions for neural commitment and differentiation. Method Enzymol 365, 327–341

Zeng, H., Horie, K., Madisen, L., Pavlova, M.N., Gragerova, G., Rohde, A.D., Schimpf, B.A., Liang, Y., Ojala, E., Kramer, F., et al. (2008). An inducible and reversible mouse genetic rescue system. PLoS genet 4, e1000069.

Zuccolo, M., Alves, A., Galy, V., Bolhy, S., Formstecher, E., Racine, V., Sibarita, J.B., Fukagawa, T., Shiekhattar, R., Yen, T., et al. (2007). The human Nup107-160 nuclear pore subcomplex contributes to proper kinetochore functions. EMBO J 26, 1853–1864.

## References cited in Supplemental Figures and Tables

Belgareh, N., Rabut, G., Bai, S.W., van Overbeek, M., Beaudouin, J., Daigle, N., Zatsepina, O.V., Pasteau, F., Labas, V., Fromont-Racine, M., et al. (2001). An evolutionarily conserved NPC subcomplex, which redistributes in part to kinetochores in mammalian cells. J Cell Biol 154, 1147–1160.

Berto, A., Yu, J., Morchoisne-Bolhy, S., Bertipaglia, C., Vallee, R., Dumont, J., Ochsenbein, F., Guerois, R., and Doye, V. (2018). Disentangling the molecular determinants for Cenp-F localization to nuclear pores and kinetochores. EMBO Rep 19, e44742.

Bolhy, S., Bouhlel, I., Dultz, E., Nayak, T., Zuccolo, M., Gatti, X., Vallee, R., Ellenberg, J., and Doye, V. (2011). A Nup133-dependent NPC-anchored network tethers centrosomes to the nuclear envelope in prophase. J Cell Biol 192, 855–871.

Festuccia, N., Owens, N., Papadopoulou, T., Gonzalez, I., Tachtsidi, A., Vandoermel-Pournin, S., Gallego, E., Gutierrez, N., Dubois, A., Cohen-Tannoudji, M., et al. (2019). Transcription factor activity and nucleosome organization in mitosis. Genome Res 29, 250–260.

Fontoura, B.M.A., Blobel, G., and Matunis, M.J. (1999). A conserved biogenesis pathway for nucleoporins: Proteolytic processing of a 186-kilodalton precursor generates Nup98 and the novel nucleoporin, Nup96. J Cell Biol 144, 1097–1112.

